# Taxane chemotherapy leads to breast cancer dormancy escape by stromal injury mediated IL-6/MAP2K signaling

**DOI:** 10.1101/2022.07.10.499472

**Authors:** Ramya Ganesan, Swati S. Bhasin, Upaasana Krishnan, Nagarjuna R. Cheemarla, Beena E. Thomas, Manoj K. Bhasin, Vikas P. Sukhatme

## Abstract

A major cause of cancer recurrence following chemotherapy is cancer dormancy escape. Taxane-based chemotherapy is standard of care in breast cancer treatment aimed at killing proliferating cancer cells. Here, we demonstrate that docetaxel injures stromal cells, which release protumor cytokines, IL-6 and G-CSF, that in turn invoke dormant cancer outgrowth both *in vitro* and *in vivo*. Single-cell transcriptomics shows a reprogramming of awakened cancer cells including several survival cues such as stemness, chemoresistance, as well as an altered tumor microenvironment with augmented pro-tumor immune signaling. IL-6 plays a role in cancer cell proliferation, whereas G-CSF mediates tumor immunosuppression. Pathways and differential expression analyses confirmed MEK as the key regulatory molecule in cancer cell outgrowth and survival. Antibody targeting of protumor cytokines (IL-6, G-CSF) or inhibition of cytokine signaling via MEK/ERK pathway using selumetinib prior to docetaxel treatment prevented cancer dormancy outgrowth suggesting a novel therapeutic strategy to prevent cancer recurrence.

## Introduction

Cancer that is loco-regional is typically treated with surgery and with neoadjuvant or adjuvant chemotherapy, radiation therapy or immunotherapy to prevent local and most importantly systemic recurrences. One mechanism for the latter is the awakening of cancer cells that are in a dormant state, which may not be eradicated by neoadjuvant or adjuvant treatments. Dormancy can be cellular (single dormant cells) or occur as a cluster wherein proliferation is balanced by cell death. Earlier studies have shown that surgery itself promotes early dormancy escape and accelerates relapse (1, 2). Recent studies have shown that inhibiting the surgery mediated injury response by administering pre-operative NSAIDs can eradicate micrometastases that exist at the time of surgery (3, 4). More recent studies have shown the role of chronic inflammation, injury response, fibrosis and autophagy inhibition in awakening, or elimination of dormant cancer cells (3, 5-8).

Chemotherapy is given in neoadjuvant and adjuvant settings, as well as standalone therapy to reduce tumor burden. However, in addition to the known toxicities of such drugs, there is an increased appreciation of other deleterious effects of such treatments. Docetaxel (T), a second generation taxane, is given to breast cancer patients as a single agent (doses ranging from 60-100 mg/m^2^) or in combination with other chemotherapeutics such as doxorubicin (A) and cytoxan (C), in cycles. Some pre-clinical studies have shown the effects of chemotherapy in inducing dormancy (9, 10), while others have focused on stress, senescence associated secretory phenotype and chemotherapy induced cell debris in cancer dormancy awakening, tumor growth and metastasis (11-14). To date the mechanistic basis for the effects of chemotherapy on cancer dormancy *in vivo* has not been elucidated.

Using a model of breast cancer dormancy, we show for the first time that taxane-based chemotherapy induces pro-inflammatory cytokine release by injured stromal cells which causes cancer dormancy escape potentiating aggressive immune evasion and cancer outgrowth. We find that inhibiting IL-6 and/or G-CSF mediated MEK signaling cascade prevents chemotherapy induced cancer dormancy escape.

## Materials and Methods

### Cell lines

D2.0R-luc-mcherry and D2.0R-FUCCI cells were obtained from Dr. Mikala Egeblad’s lab in Cold Spring Harbor Laboratory (5). 2H11 (cat # CRL-2163) and MEF (cat # CRL-2907) cells are murine endothelial and fibroblast cells lines, respectively, purchased from ATCC. All cell lines were cultured in DMEM supplemented with 10% FBS (cat # 30-2020, ATCC), 100 U/mL penicillin, 100 μg/mL streptomycin, 4.5 g/L glucose, 4mM L-Glutamine, 1 mM sodium pyruvate and 1.5 g/L sodium bicarbonate (cat# 30-2002, ATCC). All cell cultures were passaged at 80% confluency and tested negative for murine pathogens, including mycoplasma. All cells were maintained in a humidified 37 °C CO_2_ incubator.

### *In vitro* tumor stromal organoid dormancy model

D2.0R-luc-mCherry were co-cultured in reduced growth factor (RGF) basement membrane extract (BME) with murine endothelial cells (2H11) and/or murine embryonic fibroblasts (MEF) in DMEM supplemented with 2% FBS, 100 U/mL penicillin, 100 μg/mL streptomycin, 1 g/L glucose, 4mM L-Glutamine, 1 mM sodium pyruvate and 1500 mg/L sodium bicarbonate (cat # 11885084, Life Technologies Inc.). The tumor stromal organoid was incubated for 3-4 days in RGF BME coated plates to establish cancer dormancy, followed by treatment with vehicle, docetaxel, selumetinib, anti-IL-6 Ab, anti-G-CSF Ab, or combinations. At the end of treatment, cells were harvested for flow cytometry staining.

### FUCCI imaging

D2.0R-FUCCI cells, 2H11s or MEFs, were coated with nanoshuttle beads the day before spheroid formation. On the day of spheroid formations, D2.0R-FUCCI cells, 2H11 and/or MEFs were cultured as monotypic, double, or tumor stromal spheroid with D2.0R-FUCCI: 2H11: MEF in the ratio of 1:2:2 in DMEM with low glucose and 2% FBS for 3-4 days in a CO_2_ incubator. Then media was replaced with fresh media containing vehicle, docetaxel, anti-IL-6, anti-G-CSF, selumetinib or combinations. These cells were imaged using multi-channel Incucyte Live Imaging system or ECHO Revolve.

### Cell viability assay

Nano shuttle coated D2.0R cells, 2H11 cells or MEF cells were cultured for 3-4 days in DMEM + 5% FBS, seeded at 5000 cells/well in a 96-well plate to allow spheroid formation with help of a driver magnetic base. Spheroids were then treated with different doses of docetaxel (0-10 μM) in fresh culture medium. At the end of treatment, plate was equilibrated at room temperature for 30 mins and 100 μl CellTitre Glo 3D (Promega) cell viability assay reagent was added to 100 μl of cells in the well with vigorous shaking for 5 mins. Cells were incubated for 25 mins at room temperature and luminescence was recorded using Clariostar Plus microplate reader.

### Cell invasion assay

D2.0R cells were labeled with CellTracker Deep Red dye and suspended in chemotaxis buffer (DMEM containing 0.1% BSA + 2% RGF BME). CellTracker labelled D2.0R cells were mixed with 2H11 and MEF cells in the ratio of 1:2:2 at 5 ×10^4^ cells/ml in chemotaxis buffer (DMEM + 0.1% BSA+2% RGF BME). Next, 2.5 × 10^4^ cells were applied to the upper chambers of 8-μm PET Growth Factor Reduced Matrigels (24-well format), 0.5 ml chemotaxis buffer was added in the bottom well and cells were incubated for 1 day at 37 °C in a humidified 5% CO_2_ incubator. Cells were treated with 1 μM DTX and incubated at 37 °C in a humidified 5% CO_2_ incubator for 2 days. At the end of this, cells were scraped from inside the Matrigel insert, the bottom side was fixed and stained with DAPI. The peeled GFR Matrigel was carefully mounted on a glass slide. 2 separate fields of cells were counted for each invasion assay at 10x objective and expressed in terms of total number of invading cells ±S.E.

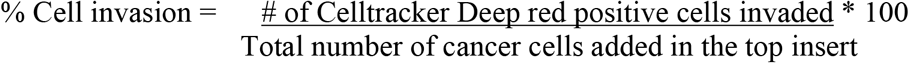

### *In vivo* tumor dormancy model

For orthotopic tumor dormancy, 5 × 10^4^ D2.0R luc-mCherry cells were injected in the 4th mammary fat pad of 6-8 weeks old immunocompetent Balb/c mice. Metastatic tumor dormancy was established by injecting D2.0R luc-mCherry cells intravenously (0.5 × 10^6^) lung metastatic dormancy in 6-8 weeks old immunocompetent Balb/c mice. Cell viability was assessed by trypan blue exclusion prior to injection and was always above 90% viability. All procedures were approved by the Emory University Institutional Animal Care and Use Committee and conformed to the Guide for the Care and Use of Laboratory Animals.

### Bioluminescence Imaging

Mice were injected with 150 mg/kg of D-Luciferin (Cat # LUCK, Gold Bio Inc.) intraperitoneally under isoflurane anesthesia. Mice were imaged using IVIS Spectrum with bioluminescence settings on the Living Image software. The bioluminescence intensity was computed as total flux (p/s) by Living Image software normalized to background bioluminescence.

### Proteomic measurements

Cytokines in the discovery sample set were measured using the BioPlex 200 MD31 Mouse Cytokine Array/Chemokine Array (Eve Technologies, Calgary, AB). Cell culture supernatant or mouse plasma was collected and shipped overnight on dry ice to Eve Technologies for analysis by BioPlex 200 MD31 multiplex immunoassay. Cytokines that were below the lower limit of quantitation were excluded from downstream analyses. Results show the standard error of mean of samples.

### Drug Administration

All drugs were administered via intraperitoneal injection except selumetinib, which was given twice daily by oral gavage. Docetaxel (USP) was diluted in 0.9% saline and administered once (day 0) and selumetinib (Selleckchem, S1008) was dissolved in DMSO/PEG/sterile PBS and administered for 8 days starting the day before chemotherapy. For cytokine ablation, mice were treated with 200 μg anti-mouse IL-6 (MP5-20F3, Bio X Cell) once on alternate days and/or 10 μg anti-mouse G-CSF (67604, R&D systems) every day for 8 days starting the day before chemotherapy. Vehicle or a rat IgG1 isotype control (HRPN, Bio X Cell), were administered to mice who did not receive the drugs or cytokine ablation.

### Flow Cytometry

For *in vitro* experiments, tumor stromal organoids (TSO) were digested with TrypLE and Accumax and passed through 40 μm strainers to obtain single cell suspension. For *in vivo* experiments, 4^th^ mammary fat pads (mfp) were collected from mice at necropsy. To prepare for flow cytometric analysis, mfp was digested in DMEM/F-12 (1:1) media containing collagenase, hyaluronidase, DNAse I, and passed through a 70 μm strainer to obtain a single cell suspension. Red blood cells were lysed using the ACK lysis buffer. Samples were then incubated with LIVE/DEAD™ Fixable Aqua Dead Cell Stain (ThermoFisher-L34957). After washing, samples were incubated in the presence of an anti CD16/32 Fc receptor-blocking antibody followed by surface staining of the following antibodies in three panels: Panel 1: APC-Cy7 CD3, FITC CD4, PerCP-Cy5.5 CD8, APC CD25 (BD 557596, 551162, 565159) (ThermoFisher 11-0042-86, 25-0441-82, CD25); Panel 2: (PE-Cf594 Ly6G, PerCP-Cy5.5 Ly6C, FITC CD11b) and Panel 3: Alexa fluor 594 anti-mCherry (ThermoFisher-M11240), Alexa fluor 647 CD34 (BD 560230), FITC Ki67 (BD 556026) with the appropriate isotype controls.

FoxP3 or Ki67 staining was performed using the eBioscience™ FoxP3/Transcription Factor Staining Buffer Set (00-5523-00), a PE FoxP3 antibody (12-5773-82) and a FITC Ki67 antibody (BD 556026). All samples were run on a Cytek Aurora flow cytometer and analyzed using FlowJo software.

### Immunohistochemistry

Tumors/mammary fat pads were excised from mice. These tissues were fixed in 10% neutral buffered formalin, embedded in paraffin, and stained with H&E. Immunohistochemical staining was performed by the Cancer Tissue and Pathology Core Lab at Winship Cancer Institute of Emory University.

### Quantification of metastatic burden and metastatic foci

The metastatic burden in lungs was evaluated using hematoxylin and eosin-stained lung sections. Metastatic burden was plotted as area of several tumor foci in the lungs, calculated as percentage of lung area.

### Single cell RNA sequencing library preparation, sequencing, and analysis

For *in vitro* experiments multi-cellular tumor stromal spheroids treated with VEH or DTX were digested with TrypLE and Accumax and passed through 40 μm strainers to obtain single cell suspension, which was processed using 10x genomics kits. For *in vivo* experiments tumors/mammary fat pads from control and docetaxel-treated mice at 45 days post-chemotherapy, were processed as described in flow cytometry above for preparing single cell suspension and downstream processing was done using the 10x genomics kits. ScRNA-seq libraries were prepared using the Next GEM Chromium single cell 3′ reagent kits V3.1 with feature barcode technology for cell surface protein (10x genomics) and sequenced using NextSeq 500 high output kits and Novaseq S4 PE100 kits (Illumina). scRNA-Seq data after standard quality control was aligned to the reference genome (mm10) using the 10x Cell Ranger pipeline. Preprocessed and filtered normalized data were subjected to unsupervised analysis using principal component analysis (PCA) (Seurat v2.0 Bioconductor package(15)) to identify principal components with significant variation that was used as input for UMAP analysis to determine overall relationship among cells. Cells with similar transcriptome profiles were clustered together, and the clusters were subsequently annotated to different cell types based on expression of specific transcripts, e.g., Endothelial cells (*Cd34+, Eng+, Pecam1+*), Fibroblasts/CAFs (*Fbn1+, Cd34+, Pdpn+*), cancer cells (*Krt18+, Krt8+*). Transcripts significantly associated with a particular cell type were identified by comparing the gene expression profile of the target cell with the rest of the cells using nonparametric Wilcox’s rank test (P < 0.01) and fold change (>1.2).

### CellChat analysis

Cell-cell communication analysis was performed using Cellchat (16) individually on the dormant and DTX treated groups and the mergeCellChat() function was used to compare them. Single cell data was input as a normalized count matrix containing the expression of all the genes present in the dataset which helped uncover the underlying pathways associated with the two groups of analyses.

### SDS-PAGE and Western blotting

To confirm the MEK activity we measured levels of MEK1/2, ERK1/2 and phospho-ERK1/2 proteins in D2.0R cancer spheroids treated with conditioned media from stromal spheroids and performed SDS-PAGE and western blot analyses specific for each of these three proteins, as well as using GAPDH as the equal protein loading control. Approximately 50 μg of protein lysate (lysis buffer composition was: 50 mM HEPES, pH 7.2, 100 μM Na_3_VO_4_, 0.1% Triton X-100 and 1 mg/ml each of protease inhibitors (aprotonin and leupeptin) was loaded per each lane of the SDS-gels that were then used for Western blot analyses. For western-blot analyses, mouse phospho-ERK1/2 (MILAN8R) IgG (eBioscience, Thermofisher) (cat. # 14-9109-82), rabbit ERK1/2 IgG (Cell Signaling Technology) (cat. # 9102S), rabbit MEK1/MEK2 IgG (SR13-07) (eBioscience, ThermoFisher) and rabbit GAPDH IgG (Cell Signaling Technology) (cat. # 97166) were utilized as primary antibodies according to the manufacturers’ recommendations. Anti-rabbit or -mouse IgG HRP antibodies were from Cell Signaling Technology (cat. # 7074 and 7076, respectively). Immunoreactivities were detected using SuperSignal West Femto chemiluminescence substrate (ECL) reagents from ThermoFisher (cat. # 34095) and Bio-Rad Chemidoc XRS+.

### Statistics

Statistical analyses were performed as described in individual figure legends. Generally, P < 0.05 was considered significant and statistical tests for *in vitro* and *in vivo* experiments were two-tailed, unless otherwise indicated. Statistics for transcripts significantly associated with a particular cell type in the single-cell RNAseq data were identified by comparing the gene expression profile of the target cell with the rest of the cells using nonparametric Wilcox’s rank test (P < 0.01) and fold change (>1.2). For flow cytometry analyses statistical significance were conducted with one-way analysis of variance (ANOVA) with Brown-Forsythe F-test followed by Dunnett’s multiple comparisons post hoc test for comparing different treatment groups, unless otherwise indicated. For *in vivo* experiments, independent t-test and one-way ANOVA with Dunnett’s posthoc comparisons were utilized. Independent t-test was utilized to evaluate significance in *in vitro* experiments with less than three treatment conditions. The Kolmogorov-Smirnov test was used to evaluate the assumption of normality of continuous variables, and no significant departures from normality were detected. When applicable, data distribution was assumed to be normal. Summary data are reported as mean ± SEM. Longitudinal luminescence intensity data were analyzed using ordinary 2-way ANOVA followed by Dunnett’s post hoc comparisons between different treatment groups. Biological replicates for each experiment are noted in figure legends. No data were excluded from the analyses. Mice were randomized from different cages and allocated to vehicle and treatment groups for all *in vivo* experiments. Immunohistochemistry images were acquired and analyzed in a blinded fashion. For all other experiments, neither randomization nor blinding was used.

## Results

### Chemotherapy breaks cancer cells out of dormancy and induces cancer proliferation and invasiveness

To investigate the effects of chemotherapy on tumor mass or cluster cancer dormancy in the tumor stromal environment, we extended a well-characterized model of cancer dormancy using D2.0R breast cancer cells (5, 6) into a physiologically relevant *in vitro* tumor stromal organoid (TSO) model comprising cancer cells, fibroblasts and endothelial cells as described in materials and methods. To establish this model, we first characterized the traditional monotypic 3D culture (D2.0R 3D) (6) and our extended TSO culture (TSO) using droplet based single-cell RNAseq analysis showing the transcriptome profiles of 3370 and 2375 individual cells from D2.0R 3D and TSO culture, respectively (Fig. 1A). A combined UMAP cluster map and heatmap show the relative abundance and transcriptome profiles of cell types in the monotypic D2.0R 3D culture and the TSO culture (Fig. 1B, S1A). To understand the transcriptome profiles of cancer cells in monotypic vs TSO culture, we subset out cancer cell clusters (i.e, D2.0R 1, D2.0R 2 and D2.0R 3) and noticed that cluster D2.0R 1 was predominantly present in the D2.0R 3D culture (monotypic), whereas cluster D2.0R 3 was predominantly present in the TSO culture. The only cluster commonly present in both the culture models was cluster D2.0R 2 (Fig. S1B). Furthermore, by differential gene expression analysis (Fig 1C) we observed that the transcriptomic profile of cancer cells in the D2.0R 3D (i.e., cluster D2.0R 1) shared very few similarities with those in the TSO culture (i.e., cluster D2.0R 3). The cluster that is common to both culture systems (i.e., cluster D2.0R 2) exhibits cell cycle and proliferation associated genes such as *Top2a, Ube2c, Mki67*, etc. These findings showed that even in monotypic 3D (D2.0R only) culture, all the cells do not undergo cellular dormancy. Some cancer cells continue to proliferate as shown by cancer cells cluster D2.0R 2, suggestive of cancer cluster dormancy rather than cellular dormancy. Furthermore, by differential gene expression analysis, we identified several dormancy-associated genes such as *Cst6, Mgp, Mme, Gas6*, etc. to be upregulated in cancer cells in the TSO culture compared to cancer cells in the monotypic culture (Table S1). Thus, after characterizing and finding cluster cancer dormancy by transcriptomics analysis in the TSO culture, we set out to study the effects of chemotherapy on dormancy.

**Figure 1.**
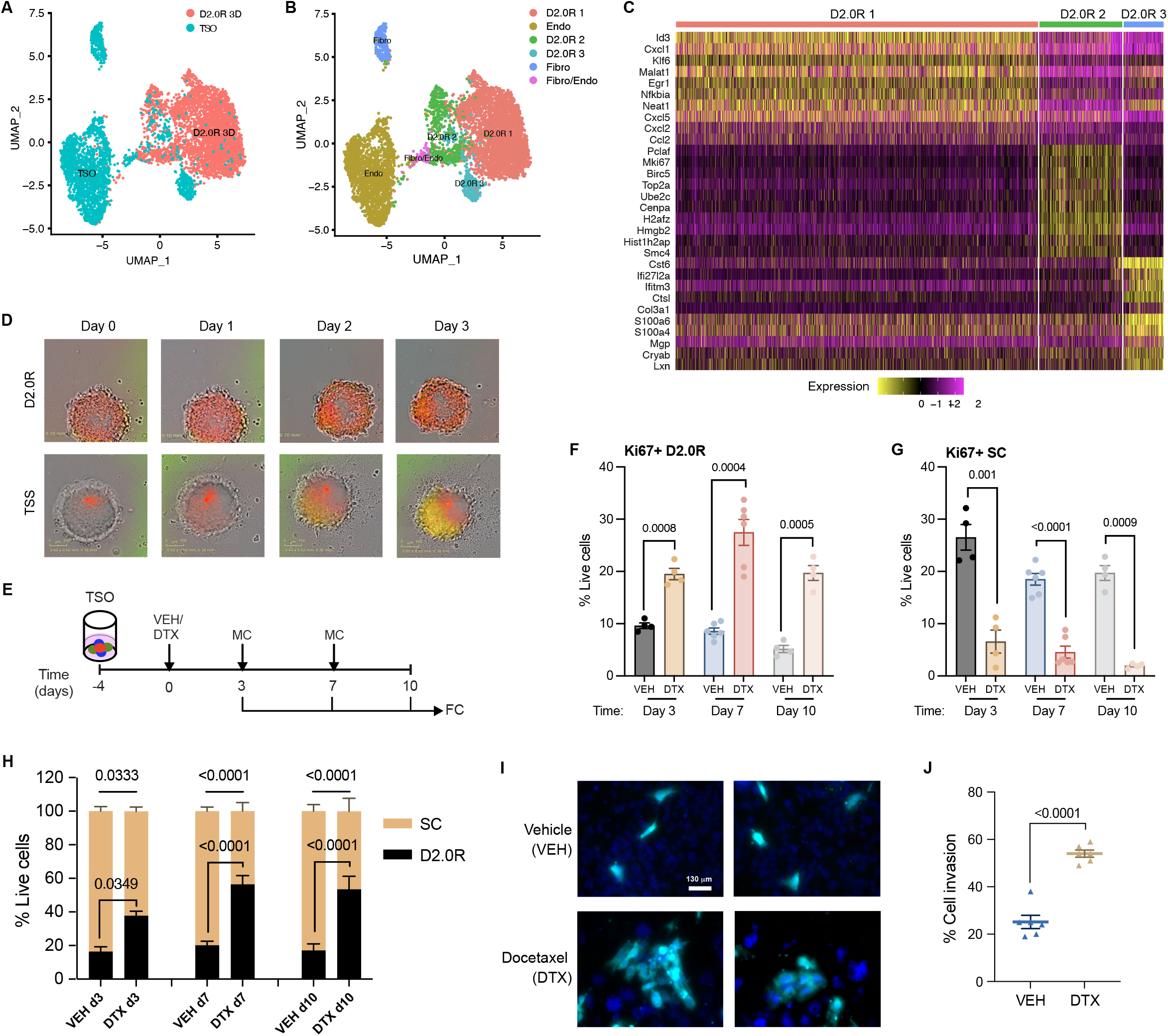
Docetaxel invokes dormancy outbreak in tumor stromal organoid and promotes invasion. (A) UMAP presentation of monotypic 3D culture (D2.0R 3D) (n=24) and tumor stromal organoids (TSO) (n=24) from scRNA-SEQ analysis. (B) UMAP presentation of major cell type clusters from scRNA-SEQ analysis in a merged dataset. (C) Heatmap showing the top 10 DEGs in each of the cancer cell clusters in a merged dataset. (D) Representative FUCCI reporter images of cancer spheroids (D2.0R) and tumor stromal spheroids/TSS (D2.0R-FUCCI: Endothelial cells (ECs): Fibroblasts (Fibro)) upon docetaxel treatment (n=3); scale 360 μm2. (E) Experimental design used to test the effects of chemotherapy on tumor stromal organoids; VEH-Vehicle, DTX-Docetaxel, MC-Media change, FC-Flow cytometry. (F-G) Expression (%) of Ki67 in (F) cancer cells (D2.0Rs) and (G) stromal cells (ECs and Fibroblasts) in tumor stromal organoid upon docetaxel (DTX) treatment compared with vehicle treated control on days 3 (n=4), 7 (n=6) and 10 (n=4) from the start of treatment. Independent t-test measurement shows statistical significance between treatment groups. (H) Stacked bar graph showing percentage of cancer cells and stromal cells out of total live cells in vehicle and docetaxel treated tumor stromal organoid on days 3 (n=4), 7 (n=6) and 10 (n=4). Independent t-test measurement shows statistical significance between treatment groups. (I) Representative images of CellTracker Deep Red dye labeled cancer cells (cyan) that invaded through the Matrigel upon vehicle or docetaxel treatment with DAPI (blue) staining for nuclei. (J) Quantification of total number of cancer cells (cyan) invaded through the matrix relative to total cells added per well (%), n=3 independent experiments in duplicates. Independent t-test measurement shows statistical significance between treatment groups.

Using physiologically relevant concentrations of docetaxel (0.01 - 10μM), we performed a dose response cell viability assay to determine the optimal concentration of docetaxel for our study. We observed that even at the highest concentration (10 μM) of docetaxel, cancer cell viability was unaffected, whereas concentrations as low as 1 μM reduced stromal cell viability significantly (Fig. S1C-E). Using D2.0R cells expressing the fluorescence ubiquitination cell cycle indicator (FUCCI) reporter, we monitored cancer dormancy or cell cycle arrest (noted by red cells) and cancer cell proliferation or cell cycle re-entry (noted by orange-yellow-green cells). We observed that when cancer spheroids (comprising cancer cells alone) are treated with vehicle (VEH) or 1 μM docetaxel (DTX), cancer cells continue to remain in a state of cell cycle arrest or dormancy (noted by red cells) (Fig 1D, top row), whereas docetaxel treatment of tumor stromal spheroids (TSS) (comprising cancer cells and stromal cells) induced cell cycle re-entry in cancer cells (noted by orange-yellowgreen cells) (Fig. 1D, bottom row). Similar results as TSS were seen when cancer cells co-cultured with either fibroblasts or endothelial cells were treated with docetaxel (data not shown). To quantitatively measure cell proliferation or tumor dormancy escape, we performed flow cytometry analysis to determine the percentage of Ki67 expressing D2.0R cells (*Flow gating* shown in Fig. S1F) on tumor stromal organoids. We observed that following docetaxel treatment there was a significant increase in the percentage of cancer cells expressing Ki67, while there was a decrease in stromal cells expressing Ki67 (Fig. 1E-G). We also noticed that the increase in proliferation corresponds to an increase in the percentage of total D2.0R cells in the tumor stromal organoid (Fig. 1H). Next, we investigated if DTX treatment impacts cancer cell invasion. We treated D2.0Rs, co-cultured with fibroblasts and endothelial cells in a trans-well insert (coated with RGF Matrigel), with DTX or VEH. We observed an increase in the number of cancer cells that invaded through the Matrigel following DTX treatment compared to VEH treatment (Fig. 1I-J). These findings suggest that chemotherapy not only causes low proliferative or slow cycling cancer cells to become highly proliferative, but also increases their invasiveness.

### Pro-inflammatory mediators from dying stromal cells invoke cancer dormancy escape

Having noticed that stromal cells succumb to chemotherapy while cancer cells remain intact, we set forth to determine if stromal injury by chemotherapy and release of secretory factors was causing cancer dormancy escape. To investigate this, we utilized multiplex cytokine assay to determine levels of 32 secretory cytokines, chemokines and growth factors in the culture supernatant. Interestingly we found that G-CSF (P= 0.0468), GM-CSF (P= 0.0007), IL-6 (P= 0.0149), KC (P= 0.0104), MIP-2 (P= 0.0079), and TNFα (P= 0.0212) were significantly elevated in the tumor stromal organoid supernatant upon docetaxel treatment (Fig. 2A). We also observed that VEGF levels were significantly decreased after chemotherapy (Fig. 2A). To identify the source of these secretory factors we subjected cancer spheroids (comprising of only cancer cells) and stromal spheroids (comprising of only stromal cells) to docetaxel treatment and found that the stromal cells were releasing these chemokines, cytokines, and growth factors (Fig. 2B-G). Based on our findings (Fig. 2A-G) and taking into consideration previous studies showing the protumor effects of IL-6 and G-CSF (17), we further investigated the role of these pro-inflammatory mediators in cancer dormancy escape. We found that neutralizing antibodies against IL-6 and G-CSF starting two days prior to docetaxel treatment significantly inhibited chemotherapy induced cancer cell proliferation (Fig. 2H). We also quantified this by flow cytometry staining and analysis (Fig. 2I-J). We observed that treatment with neutralizing antibodies alone against IL-6 or G-CSF or both together did not affect cell proliferation (both cancer and stromal cells). However, treatment with neutralizing antibodies starting two days prior to chemotherapy prevented cancer cells from becoming proliferative (Fig. 2I), while stromal cell killing by chemotherapy was not reverted by cytokine blockade (Fig. 2J). These findings confirm that IL-6 and G-CSF are critically important mediators in causing cancer dormancy escape.

**Figure 2.**
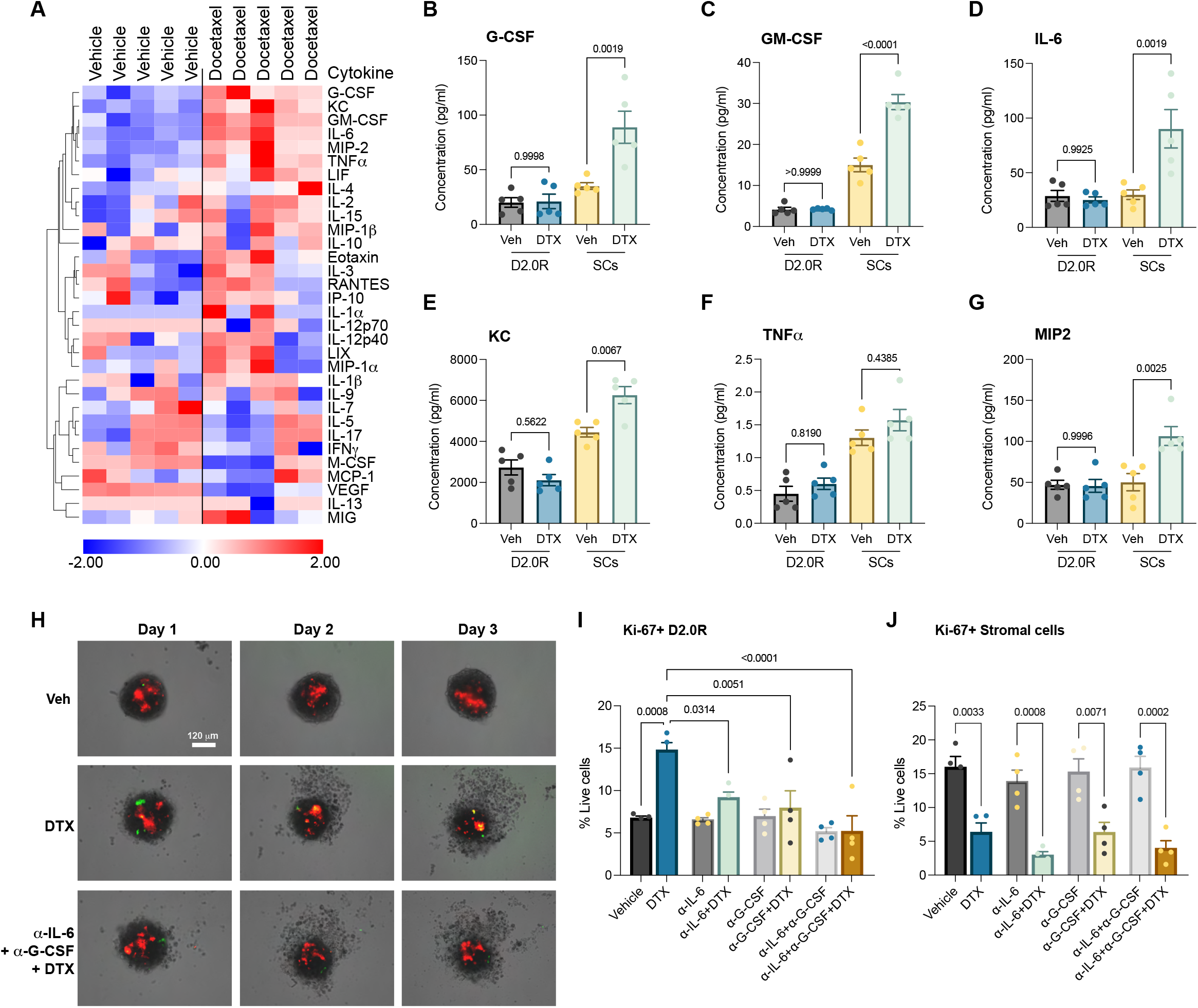
Multiplex secretory protein analysis shows chemotherapy injures stromal cells releasing proinflammatory mediators that invoke cancer dormancy awakening. (A) 32-plex cytokine analysis of tumor stromal organoid culture supernatant upon vehicle or docetaxel treatment (n=5). (B-G) Concentrations of G-CSF (B), GM-CSF (C), IL-6 (D), KC (E), TNFα (F), MIP2 (G) in culture supernatants of cancer spheroids or stromal spheroids treated with vehicle or docetaxel (n=5, each). One-way ANOVA measurement with posthoc Tukey’s multiple comparison test for flow cytometric analysis shows statistical significance between respective treatment groups. (H) Representative FUCCI reporter images of tumor stromal spheroids (D2.0R: ECs: Fibro) upon vehicle (Veh), docetaxel (DTX) or cytokine blockade with docetaxel treatment (anti-IL-6+anti-G-CSF+DTX) (n=3); scale 120 μm. (I-J) Expression (%) of Ki67 in (I) D2.0Rs and (J) stromal cells in tumor stromal organoid upon docetaxel treatment compared to vehicle treated control or cytokine blockade with or without docetaxel treatment (n=4), with 6 replicated per experiment. One-way ANOVA measurement with posthoc Dunnett’s multiple comparison test for flow cytometric analysis shows statistical significance between respective treatment groups.

### Transcriptomic analysis revealed involvement of MEK signaling in dormancy awakening

To further investigate molecular signaling and pathways involved in cancer dormancy escape induced by chemotherapy, we performed single cell transcriptomic analysis. We utilized droplet-based single cell RNA sequencing (scRNA-Seq) assay to obtain single-cell resolution transcriptomes of tumor stromal organoids. The transcriptomic profiles of 1738 and 1293 individual cells derived from vehicle and docetaxel-treated organoids, respectively revealed that docetaxel modulated the tumor stromal landscape by altering the relative abundance (Fig. 3A) and transcriptome profiles of cancer cells, fibroblasts, and endothelial cells (Fig. S2A-B). We observed by split Uniform Manifold Approximation and Projection (UMAP) cluster analysis that the relative abundance of cancer cells was increased in the docetaxel treated samples (Fold-Change = 2.1), while that of endothelial cells and fibroblasts was decreased (Fig. 3A). The differential expression analysis identified 1600 genes were differentially regulated in docetaxel treatment relative to vehicle control, including 734 overexpressed and 866 reduced genes (p-value <0.05) (Fig. S2B). Fig. 3B shows the top 20 upregulated genes in DTX and VEH organoid datasets, wherein genes associated with autophagy induced dormancy (*Col1a1, Stmn1*) and tumor suppression (*Tpi2, Tpm2*) were enriched in VEH group (18-20), while genes associated with chemoresistance (*Fth1, Hmox1*), breast cancer tumorigenesis (*Lcn2*), cancer proliferation and invasiveness (*Cxcl5, Spp1*) were enriched in DTX group (21-25). Ingenuity pathways analysis (IPA) of pathways associated with DTX treatment enriched transcripts demonstrated activation (z-score > 1) of multiple pathways related to cancer pathogenesis, cytokine signaling, cell proliferation and anti-apoptosis, etc. (Fig. 3C). After sub-setting cancer cells (clusters 2 and 5) alone and comparing docetaxel treated to vehicle control, we found 86 genes were differentially regulated with 30 overexpressed and 56 downregulated (p-value <0.05) in the docetaxel treated D2.0R cells (Fig. 3D). The most significantly enriched genes in D2.0Rs upon docetaxel treatment were known protumor genes such as *Fth1, Lcn2, Cxcl5, Tubb5, Tuba1a, Tubb4b* and *Hmox1*, which have been implicated in chemo-resistance (21, 26, 27), breast cancer tumorigenesis (23), invasiveness, metastatic colonization (24) and cancer cell survival (22). IPA revealed several upstream activated regulatory molecules in cancer cells including *Hras, Myc, Mek*, etc. Interestingly, several *Mek* (activation z-score= 2.2872, p-value= 1.20 ×10^−16^) related genes were seen to be upregulated in cancer cells as shown in Fig. 3E, which play a role in governing cell proliferation.

**Figure 3.**
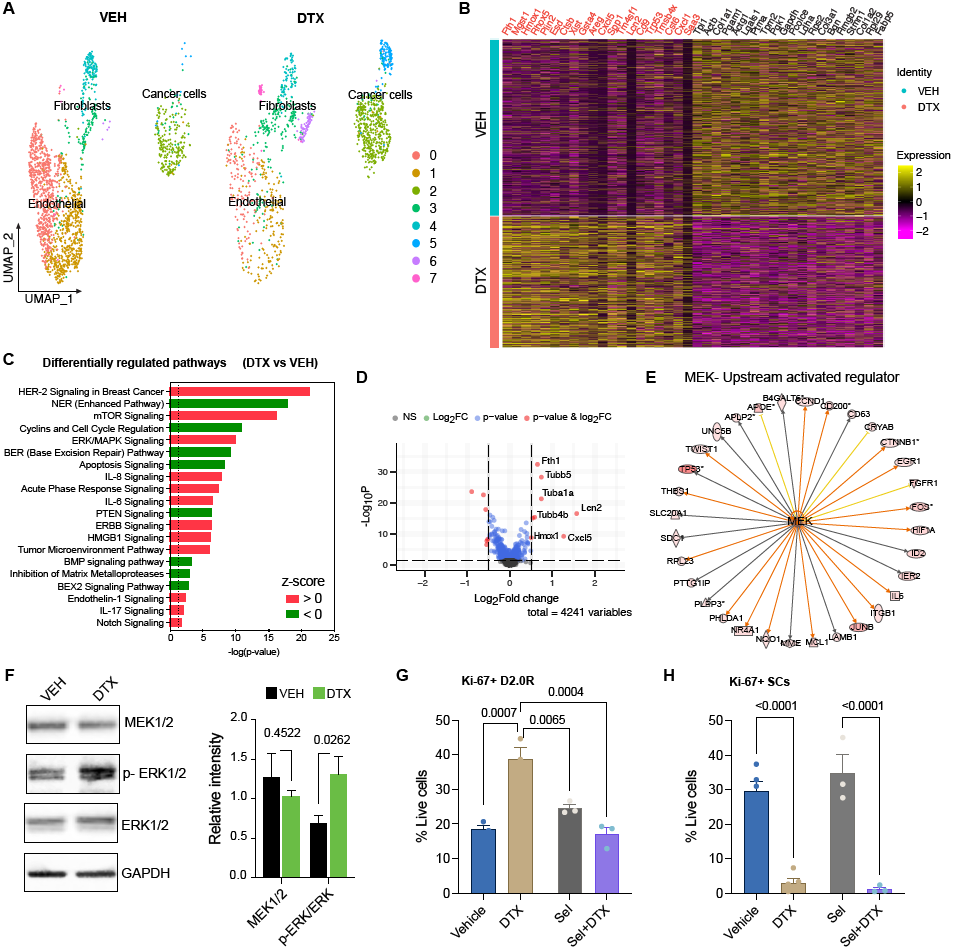
Single-cell transcriptomics revealed MEK signaling in docetaxel mediated dormancy awakening. (A) UMAP presentation of major cell type clusters from scRNA-SEQ analysis of vehicle (VEH) (n=24) and docetaxel (DTX) (n=36) treated tumor stromal organoids. (B) Heatmap showing top 20 enriched genes in DTX and VEH datasets, respectively. (C) Pathways enrichment upon docetaxel treatment compared with vehicle controls based on 1600 differentially expressed genes. Significance for enrichment is calculated based on Fisher’s exact test for each pathway are indicated on the x-axis (-log p-value); Color red or green indicates positive or negative z-score, respectively. (D) Volcano plot showing significantly differentially expressed protein-coding genes in cancer cells (D2.0R) based on RNA-seq of tumor stromal organoids from docetaxel treated compared with vehicle treated controls. Transcripts with absolute FC > 0.5 and adjusted P value < 0.05 are highlighted in red. (E) Targets of MEK enriched in cancer cells. Colored lines indicate relationships between nodes, with orange lines showing enhancement and gold lines showing inhibition of a DEG by MEK. (F) Representative western blot image showing MEK, p-ERK, ERK and GAPDH (loading control) and quantification of band intensity relative to loading control (n=3). Independent t-test measurement of band intensities of proteins shows statistical significance between docetaxel and vehicle treatment. (G-H) Flow cytometric analysis of proliferating (Ki67+) cancer cells (G) and stromal cells (H) upon vehicle (n=5), DTX (n=5), selumetinib (n=3) +/-docetaxel (n=3) treatments. One-way ANOVA measurement with posthoc Dunnett’s multiple comparison test for flow cytometric analysis shows statistical significance between respective treatment groups.

To confirm the source of *Il6* and *Csf3*, we looked at their gene expression in the tumor stromal organoids and noticed *Il6* gene expression in clusters D2.0R 2 and Fibro/CAFs 1, while *Csf3* gene expression was only seen in Fibro/CAFs 1 (Fig. S2C). Further analysis by treatment revealed that both *Il6* and *Csf3* gene expression increased in Fibro/CAFs 1 after chemotherapy, while there was a marginal decrease in expression of *Il6* in cancer cells (D2.0R 2) with no *Csf3* expression (Fig. S2D). It is widely known that MEK/MAPKK is a signaling mediator in the cytokine signaling pathway. Taken together, these findings are in accordance with the protein analysis data from multiplex cytokine assay (Fig. 2) implicating a role of stromal injury mediated paracrine cytokine signaling in cancer cell proliferation via MEK or the ERK/MAPK pathway. To confirm this, we treated cancer spheroids cultured in RGF Matrigel with conditioned media (CM) from stromal cells treated with vehicle or chemotherapy. As expected, we found an increase in MEK activity in cancer cells evidenced by no change in MEK or total ERK1/2 levels, but a robust increase in phosphorylated ERK1/2 protein levels in D2.0R spheroids upon chemo-CM treatment (Fig. 3F). Furthermore, to confirm the role of MEK in chemo-mediated dormancy awakening, we treated the tumor stromal organoid with selumetinib, a MEK1/2 inhibitor. We found that treatment with selumetinib starting one day prior to chemotherapy prevented docetaxel induced cancer dormancy escape but did not affect stromal cell killing, as shown by a significant decrease in the cancer cell proliferation (i.e., fewer Ki67+ D2.0Rs) and no rescuing effect of stromal cell proliferation (Fig. 3G-H). These findings confirmed the role of MEK/ERK signaling in docetaxel mediated dormancy awakening and indicate that MEK blockade prevents cancer dormancy outgrowth.

### Docetaxel invokes cancer dormancy escape *in vivo*

To determine whether chemotherapy could directly awaken disseminated and/or dormant cancer cells *in vivo*, we investigated both orthotopic (primary tumor) and metastatic dormancy using a syngeneic murine breast cancer model. To establish orthotopic dormancy, we injected luciferase-mCherry expressing D2.0R cells in the mammary fat pad and tracked cancer cells *in vivo* by bioluminescence imaging (Fig. 4A). We allowed the total luminescent flux to reach a steady basal level with no increasing trend. For our study, we considered the state of steady basal luminescence for at least 2-3 consecutive reads as dormancy, which was reached around 30-40 days post-inoculation (Fig S3A). Once dormancy was established, we treated mice with vehicle or docetaxel and continued measuring cancer luminescence intensity. We observed that following a single dose of chemotherapy, the luminescence intensity increased significantly over time compared to vehicle treated mice (Fig. 4B-C). We also confirmed this by H&E staining of the mammary fat pad at the end of study, wherein a single dose of docetaxel resulted in an increase in the size of tumor lesion in the mammary tissue compared to the vehicle treated control mice (Fig. 4D). A recent review on cancer dormancy has highlighted that one way of targeting dormant cells is by awakening them and then killing the awakened/proliferating cancer cells (28). Chemotherapy often preferentially targets proliferating cancer cells and is given for several cycles. To investigate if cancer cells awakened from dormancy by a single dose of chemotherapy can be killed by subsequent cycles of chemotherapy, we administered chemotherapy for 3 cycles. We found cancer cells awakened from dormancy after one dose of docetaxel did not succumb to subsequent cycles of chemotherapy as evidenced by lack of response after the third cycle of treatment (Fig. S3B-C).

**Figure 4.**
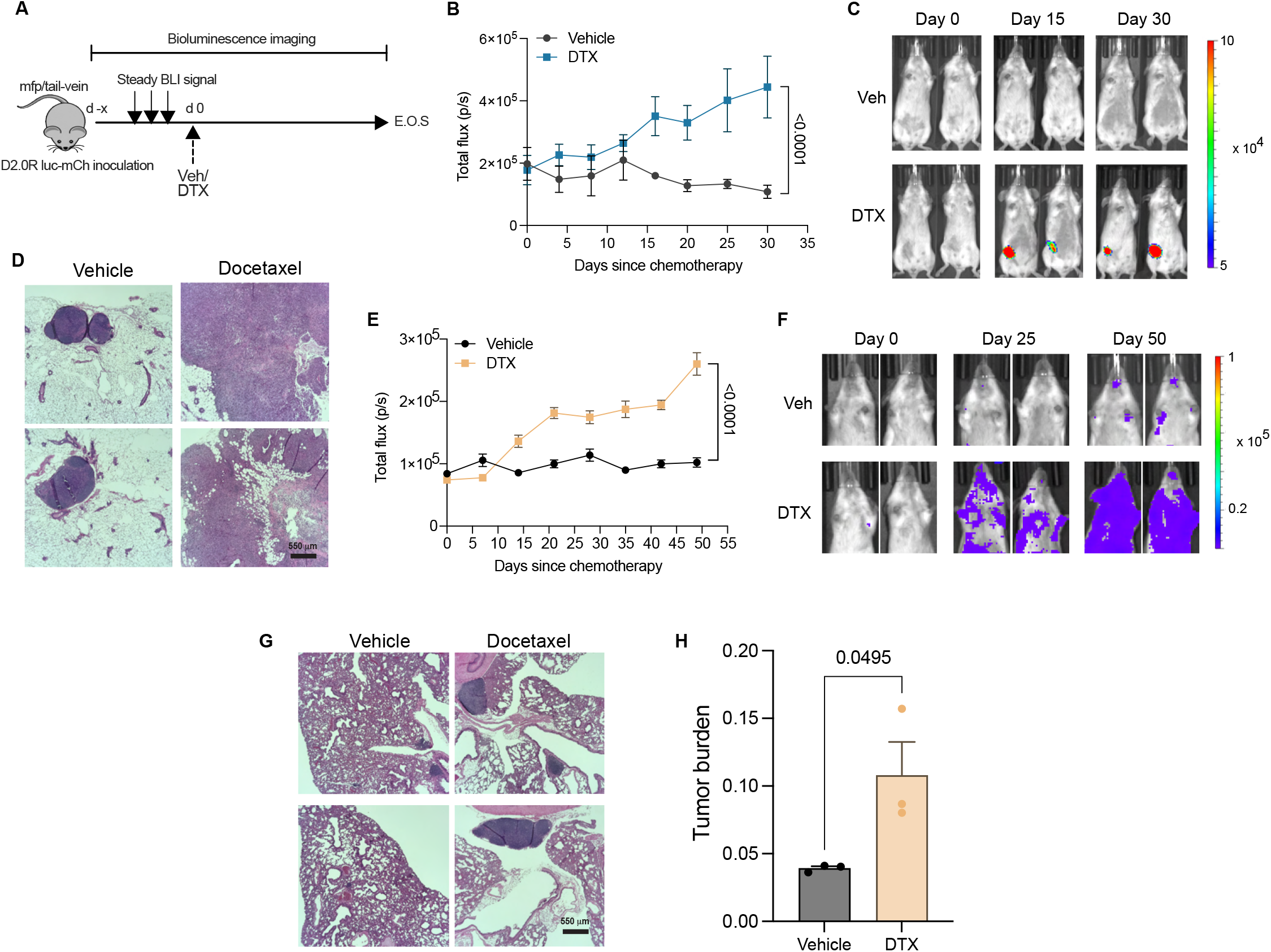
In vivo breast cancer dormancy and docetaxel mediated dormancy outgrowth. (A) Schematic showing mouse model of primary (breast) or metastatic (lung) dormancy by injection of D2.0R luc-mCherry cells in the 4th right inguinal mammary fat pad (mfp) or tail-vein, respectively; days since chemotherapy (d), docetaxel 8 mg/kg (DTX), end of study (E.O.S) and vehicle (Veh). (B-C) Bioluminescence flux kinetics (B) and representative BLI images (C) of D2.0R luc-mCherry tumor growth in the mfp of mice in docetaxel treated group (n=9) compared with vehicle treated controls (n=7). Two-way mixed ANOVA with posthoc Dunnett’s multiple comparisons test shows statistical significance between treatment groups. (D) Representative gross images of mammary fat pads (4th right inguinal), hematoxylin and eosin (H&E) staining are shown from Vehicle and Docetaxel treated mice; scale bars, 550 μm. (E-F) Bioluminescence flux kinetics (E) and representative BLI images (F) of D2.0R luc-mCherry tumor growth in the lungs of mice in docetaxel treated group (n=5) compared with vehicle treated controls (n=4). Two-way mixed ANOVA with posthoc Dunnett’s multiple comparisons test shows statistical significance between treatment groups. (G-H) Representative gross images of mouse lungs (G), hematoxylin and eosin (H&E) staining, and tumor burden quantification (H) are shown from Vehicle and Docetaxel treated mice; scale bars, 550 μm. Independent t-test measurement shows statistical significance between treatment groups.

Previous studies have shown that acute inflammation induced NET formation or fibrosis can invoke breast cancer metastatic dormancy awakening in the lungs (5, 6). To evaluate the role of chemotherapy in metastatic dormancy awakening, we employed the lung metastatic dormancy model (5) by tail-vein injection of D2.0R cells expressing luciferase-mCherry followed by vehicle or docetaxel treatment (Fig. 4A). As expected, we observed an increase in the luminescent flux following docetaxel administration (Fig. 4E-F). We also confirmed metastatic dormancy awakening upon docetaxel treatment by H&E staining of the lungs as evidenced by an increase in the lung metastatic burden (Fig. 4G-H). Thus, our findings demonstrate that chemotherapy invoked both primary/orthotopic and metastatic cancer dormancy escape in a mouse model of breast cancer.

### Chemotherapy induced systemic response and altered tumor immune landscape

Having confirmed that chemotherapy awakens dormant cancer cells irrespective of the site of dormancy, we investigated if systemic secretory factors played a role in cancer dormancy awakening. For this, we utilized multiplex cytokine assay to determine levels of 32 cell secretory factors and found that IL-6 (P= 0.0043) and G-CSF (P= 0.0003) levels were robustly increased in murine plasma upon docetaxel treatment compared to vehicle treated controls (Fig. 5A), in agreement with our *in vitro* data (Fig. 2A). Studies have shown that conventional chemotherapy alters the tumor immune landscape (29), while IL-6 and G-CSF induce neutrophil infiltration, modulate effector T cells and promote tumor progression (17, 30). To further evaluate the effects of these secretory factors on cancer cells and leukocyte infiltration in the tumor, we performed flow cytometry staining and analysis (Fig. 5B-J). We observed an altered tumor-immune landscape upon docetaxel treatment. We found that chemotherapy increased cancer cell proliferation (Ki67^+^) (Fig. 5B) and resulted in an increase in immunosuppressive myeloid cells like neutrophils, MDSCs and M2 macrophages, with no changes observed in monocytes or M1 macrophages (5C-H). In the lymphoid compartment, we observed an increase in regulatory T cells (Tregs), decrease in anti-tumor CD8+ T cells in the mammary tissue with little to no change in CD4+ T cells (Fig. 5I-K). We also observed an overall immunosuppressive signature in the tumor as shown by a decrease in the ratio of M1:M2 macrophages and CTL:Tregs (Supp Fig. S4A-B). Furthermore, by leveraging single cell transcriptomics analysis of the tumor/mammary tissue from mice treated with vehicle or docetaxel (Fig. 5L and S5A-B), we revealed an altered tumor immune microenvironment confirming our flow cytometry data. We noticed an increase in the relative abundance of neutrophils, monocytes/macrophages, γδT cells, Tregs, CD14+ regulatory DCs, NK/NKT cells and a decrease in CD4^+^ and CD8^+^T cells, B cells and effector DC populations (migratory DCs and pDCs) upon docetaxel treatment (Fig. 5M). Diving deeper, we investigated whether these tumor immune infiltrates were pro- or anti-tumor. Markers of immunosuppression, alternative macrophage activation, pro-angiogenic, hypoxia-related and inhibitory molecules gene signature(31) including Ccl5/RANTES, Ccl8/MCP-2, Cxcl1/CXCL1, Cxcl2/CXCL2, Mmp9/MMP9, Vegfa/VEGF, Tgfb1/TGF-β1, Tnf/TNF-α, Fn1/FN (Fibronectin), Spp1/OPN (Osteopontin), Hilpda/HIG, Hmox1/HO-1, Pdcd1/PD-1, Cd274/PD-L1, Lag3/LAG-3 and Havcr2/TIM-3, were expressed by these immune cells. Comparative analysis between docetaxel treatment vs. vehicle control of the pro-tumor gene signature in immune cells demonstrated an upregulation of this immunosuppressive gene signature upon docetaxel treatment (Fig. 5N). These findings confirmed that docetaxel induces systemic release of IL-6, G-CSF and altered tumor microenvironment with augmented pro-tumor immune infiltrates.

**Figure 5.**
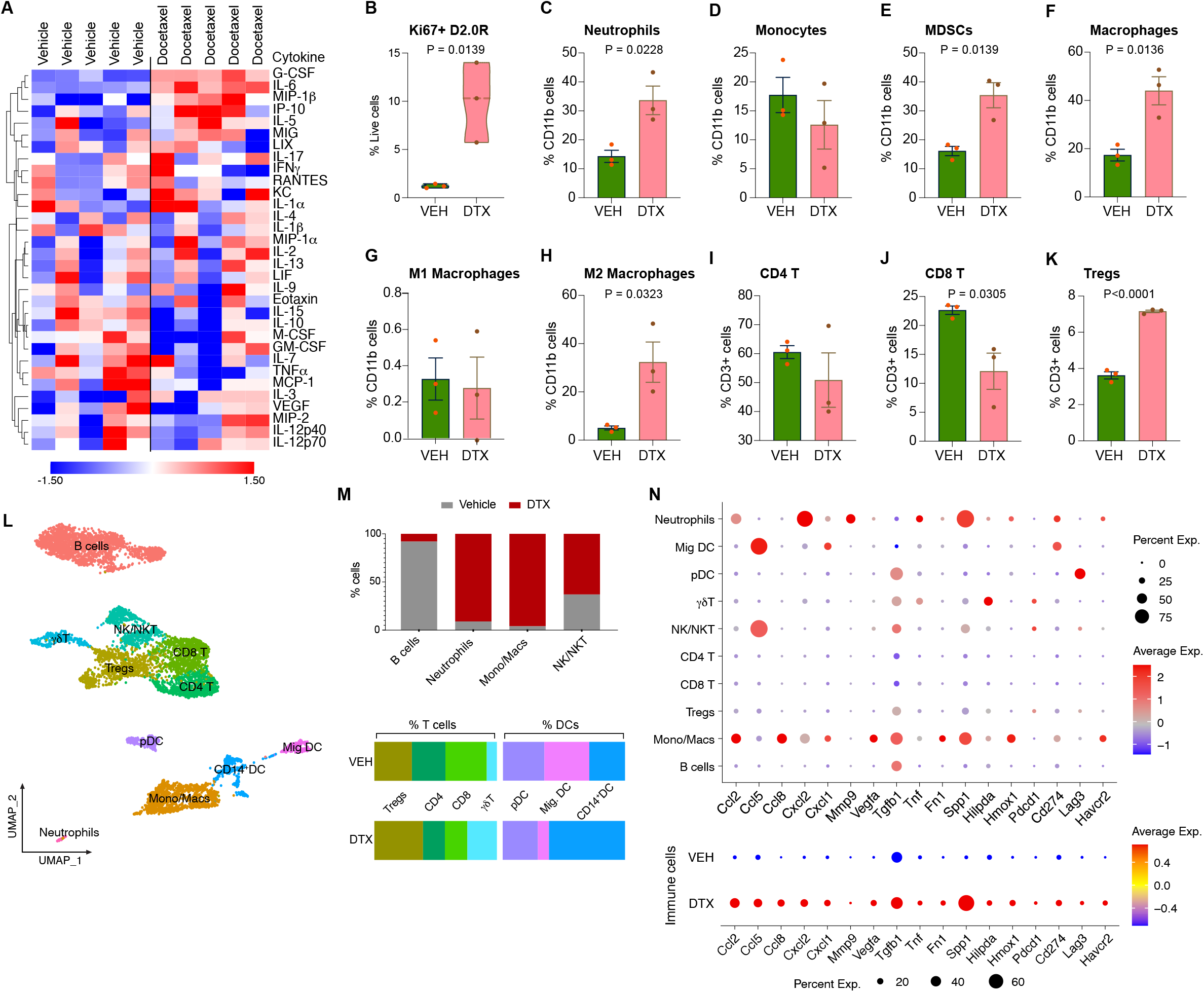
Chemotherapy induced systemic response and altered tumor immune landscape assessment using single cell transcriptomics, secretory protein, and flow cytometry analyses. (A) 32-plex cytokine analysis of plasma from mice subject to vehicle or docetaxel treatment (n=5). (B) Expression (%) of Ki67 in D2.0Rs in the mammary tumor tissue of DTX compared to VEH-treated mice. (C-K) Flow cytometric analysis showing tumor immune infiltrates including (C) neutrophils, (D) monocytes, (E) MDSCs, (F) total macrophages, (G) M1 macrophages, (H) M2 macrophages, (I) CD4 T cells, (J) CD8 T cells and (K) Tregs comparing DTX treatment with VEH control (n=3, each). Independent t-test measurement shows statistical significance between treatment groups. (L) UMAP showing immune cells in the tumor tissue of VEH and DTX datasets merged (n=2, each). (M) Percentage of cells from VEH control and DTX-treated mice, per cluster for immune cells. (N) Dot plots of selected markers from merged samples (top) and grouped by treatment, VEH and DTX (bottom). Dot size indicates the proportion of cells in each cluster expressing a gene and color shading indicates the relative level of gene expression.

### Single cell RNA-sequencing (ScRNA-seq) revealed stromal injury response invokes cancer dormancy outgrowth

scRNA-Seq of the mouse mammary tumor/tissue yielded transcriptomic profiles of 6241 and 5587 individual cells from mice treated with vehicle and docetaxel, respectively. Split UMAP cluster maps revealed altered tumor microenvironment (TME) with changes in relative abundance and transcriptome profiles of cancer cells, stromal cells, and immune infiltrates (Fig 6A; Fig. S5A). The analysis depicted the relative abundance of cancer cells and non-immune stromal cells was significantly altered with increased cancer cells and decreased fibroblasts/CAFs, cancer associated fibroblasts (CAFs), fibrocytes, adipocytes/mammary epithelial cells (Adipo/MEpCs) upon docetaxel treatment compared to vehicle treated control mammary tumor (Fig. 6A, B). Interestingly, while there was only one cancer cell cluster in vehicle-treated dormant tumors (Cancer cells 1), there were two cancer cell clusters (Cancer cells 1 and Cancer cells 2) in DTX treated tumors. A comparative analysis of cell-cell communication based on ligand-receptor expression in the mammary tissue/tumor of Vehicle and Docetaxel treated mice was performed using CellChat(16), which quantitatively measures the propensity of cell types to function as a sender or receiver for several major signaling pathways. The analysis showed cancer cell clusters were highly networking in the DTX dataset (shown by blue hotspots), whereas stromal cells were highly networking clusters in VEH dataset (shown by red hotspots) (Fig. 6C; Fig. S5C).

**Figure 6.**
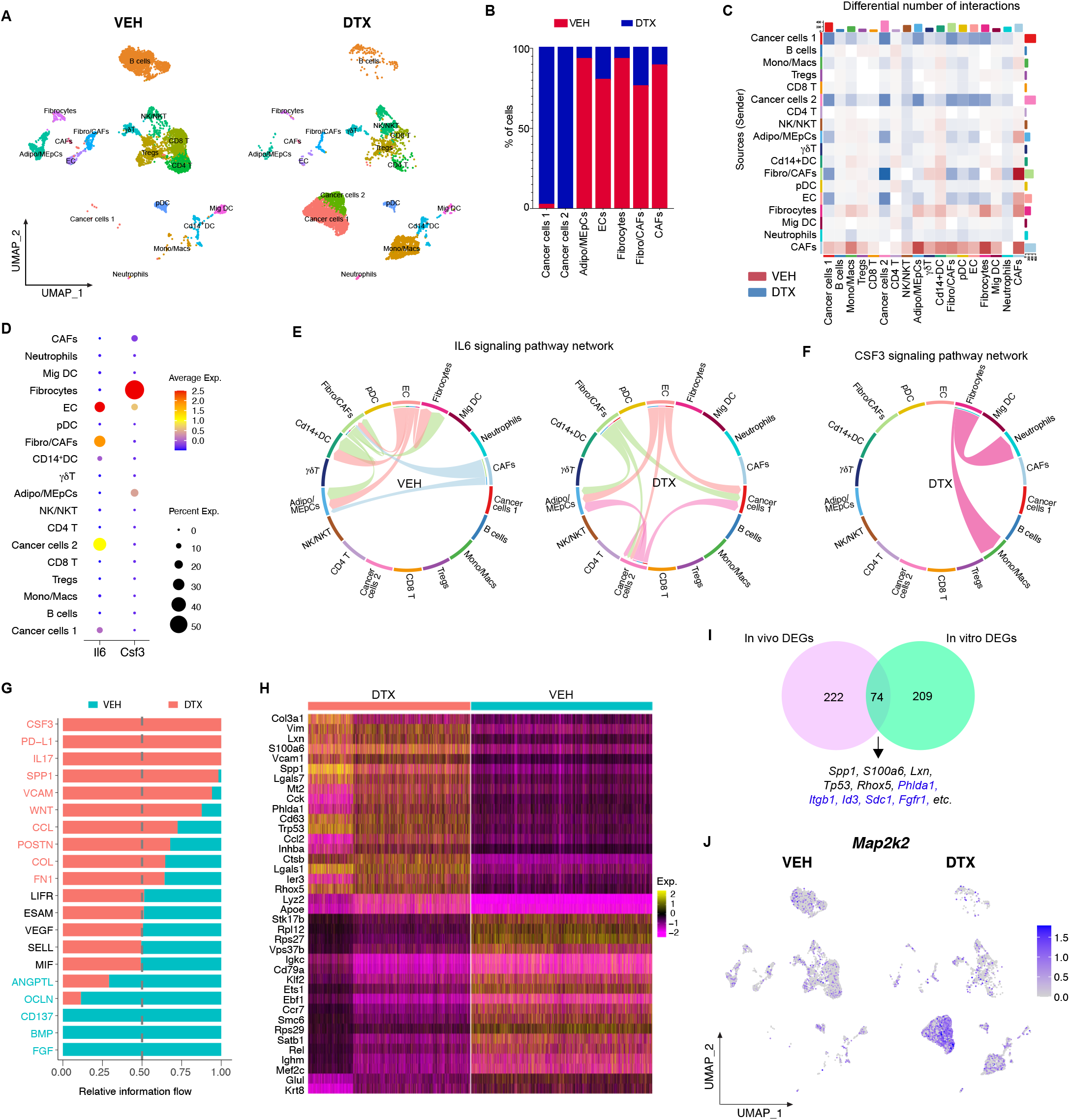
Single-cell RNA sequencing analysis of mouse mammary tumors. (A) Split UMAP presentation of major cell types and associated clusters in vehicle (VEH) and docetaxel (DTX) treated murine mammary fat pad/tumor (n=2, each treatment group). (B) Percentage of cells from vehicle control and docetaxel-treated mice, per cluster for cancer cells and non-immune stromal cells. (C) Heatmap of differential interactions between the different cell types in VEH and DTX treated groups: Rows and columns represent source and target clusters, respectively; Gradience in the color shows lowest (light) to highest (dark) number of networks; red gradient-Vehicle and blue gradient-DTX. Bar plots on the right and top of the heatmap represent the total outgoing and incoming interaction scores, respectively. (D) Dot plot showing cluster wise expression of *Il6* (IL-6) and *Csf3* (G-CSF) by clusters. (E) Chord connections between cell types shows *Il6* signaling network in Vehicle and Docetaxel datasets, respectively. (F) Chord connections between cell types showing *Csf3* signaling network in DTX dataset. (G) Quantitatively comparing the information flow of each signaling pathway between cells from VEH and DTX treated samples. The overall information flow of a signaling network is calculated by summarizing all the communication scores in that network. (H) Heatmap showing top 20 enriched genes between cells from DTX and VEH treated samples, respectively. (I) Venn showing cancer cluster DEGs in vivo (pink) and in vitro (turquoise) with overlapping genes at the intersection. (J) Split feature plot showing *Map2k* (MEK) expression in DTX vs VEH treated samples.

Having found that IL-6 and G-CSF levels were increased in plasma after DTX treatment (Fig. 5A), we asked whether the molecular signaling mechanisms were similar *in vivo* and *in vitro* (Fig. 2-3). To investigate the molecular signaling invoked by IL-6 and G-CSF upon docetaxel treatment, we first identified the source of IL-6 and G-CSF in the TME using single cell transcriptomics data. We found that the non-immune stromal cells are the primary source of *Il6* (*IL-6*) and *Csf3* (*G-CSF*) in the TME (Fig. 6D). Interestingly, we found that the cancer cells (Cancer cells 2) that propagated after DTX treatment also expressed *Il6* gene (Fig. 6D). A closer look at the IL-6 signaling pathway network revealed a shift in its role in the dormant VEH treated versus proliferating DTX treated TME (Fig. 6E). In the dormant TME, IL-6 signaling was constrained to stromal cells such as ECs and fibroblasts which are seen to interact among themselves while in the DTX treated tumor tissue, IL-6 activity occurs primarily between the cancer cells and the ECs, fibroblasts characterizing the emergence of complex interactions between cancer cells and stromal cells upon DTX treatment. IL-6 responsive genes such as *Il-17* and *Spp1(32, 33)* showed elevated signaling with active communication between cancer cells and immune and non-immune stromal cells in the tumor tissue upon DTX treatment (Fig. S5D-E). The CSF3 pathway was enriched in the DTX treated tumor tissue defined by fibrocytes interacting with neutrophils and M2 macrophages while this phenomenon was absent in the dormant state (VEH treated) (Fig. 6F). Signaling networks such as VCAM, WNT, FN1 etc. known to be associated with dormancy escape were enriched in the DTX treated tumor tissue, while BMP and FGF signaling networks, which are known to play a role in cancer dormancy (34, 35), were enriched in VEH treated dormant/control tumor tissue. Therefore, cell communication analysis provided a comprehensive overview of the transformative cellular signaling in the TME upon treatment with DTX.

We identified the top 20 enriched genes in DTX and VEH treated mammary tumor tissue, respectively and found that *Col3a1, Spp1, Vim, S100a6*, etc., which are genes associated with cancer cell proliferation, invasion, ECM remodeling and tumor progression were enriched upon DTX treatment confirming a deleterious role of DTX *in vivo* (Fig. 6H). Having observed that the molecular signaling *in vitro* and *in vivo* were similar, we asked if the cancer cells also behave similarly. For this, we performed comparison analysis between gene expression profiles of cancer cells clusters from *in vitro* and *in vivo* scRNA-Seq datasets. The resultant Venn-diagram reveals the intersection of 74 enriched genes (Fig. 6I). Of these 74 commonly enriched genes, we found several genes (highlighted in blue) that were previously shown by IPA to be associated with MEK signaling (Fig. 3E). Thus, to confirm this, we focused on the expression of *Map2k2* (*MEK*) in the mammary tumor tissue. Interestingly, we found that MEK expression was robustly increased in the cancer cell clusters upon docetaxel treatment (feature plot in Fig. 6J). We also confirmed this by IPA, wherein we found ERK/MAPK signaling to be enriched in cancer cells (Fig. S5F). Hence, these findings confirmed that docetaxel alters IL-6 and G-CSF signaling by stromal cells *in vivo* augmenting MAP2K signaling in cancer cells, thus likely awakening cancer dormancy and promoting cell proliferation.

### IL-6 and/or G-CSF signaling inhibition prevents docetaxel induced breast cancer dormancy escape *in vivo*

After confirming the systemic release of secretory factors (IL-6 and G-CSF), their molecular signaling and alterations to the tumor immune landscape, we next investigated whether inhibiting them would prevent cancer dormancy escape. For this, we performed cytokine ablation (using neutralizing antibodies) or inhibition of the cytokine signaling mediator MEK (using selumetinib) prior to chemotherapy and studied *in vivo* dormancy vs awakening (Fig. 7A). After establishing dormancy, mice were treated with neutralizing antibodies against IL-6 and/or G-CSF, isotype control or selumetinib, starting the day before chemotherapy. We observed that cytokine ablation with neutralizing antibodies against IL-6 or G-CSF or both significantly inhibited chemotherapy induced dormancy awakening as shown by *in vivo* luminescence imaging (Figs. 7B-C). Similarly, we found that selumetinib shown by transcriptomic analysis to mediate the effects of proinflammatory factors in cancer dormancy outgrowth also inhibited the deleterious effects of chemotherapy as evidenced by a significant decrease in luminescent photon flux compared to the docetaxel treated mice and no significant difference between selumetinib treated mice and vehicle controls (Figs. 7B-C). Next, we set out to determine the effects of cytokine ablation or MEK inhibition on the tumor immune landscape. For this, we performed multicolor flow cytometry analysis (Fig. 7D-K) and found that in the presence of neutralizing antibodies against IL-6 and/or G-CSF, or inhibition of MEK, the percentage of Ki67+ cancer cells was significantly reduced suggesting rescue from DTX induced dormancy outgrowth (Fig. 7D). We also observed a decrease in the pro-tumor immune infiltrates, such as neutrophils, MDSCs and Tregs upon cytokine ablation or MEK inhibition, while an increase in cytotoxic CD8+ T cells (Fig. 7E-I) was only seen with selumetinib treatment. We noticed a robust reversal of immunosuppressive signatures shown by an increase in the ratios of M1:M2 TAMs and CD8:Tregs, which are important in predicting OS, pCR or RFS (36, 37), with selumetinib treatment starting prior to DTX treatment (Fig. 7J-K). These findings confirm the causal role of IL-6 or G-CSF mediated MEK signaling in cancer dormancy awakening and tumor immunosuppression. Thus, inhibiting the inflammatory mediators (IL-6, G-CSF) or downstream MEK signaling prevented chemotherapy (DTX) induced breast cancer dormancy awakening and reduced tumor immunosuppression.

**Figure 7.**
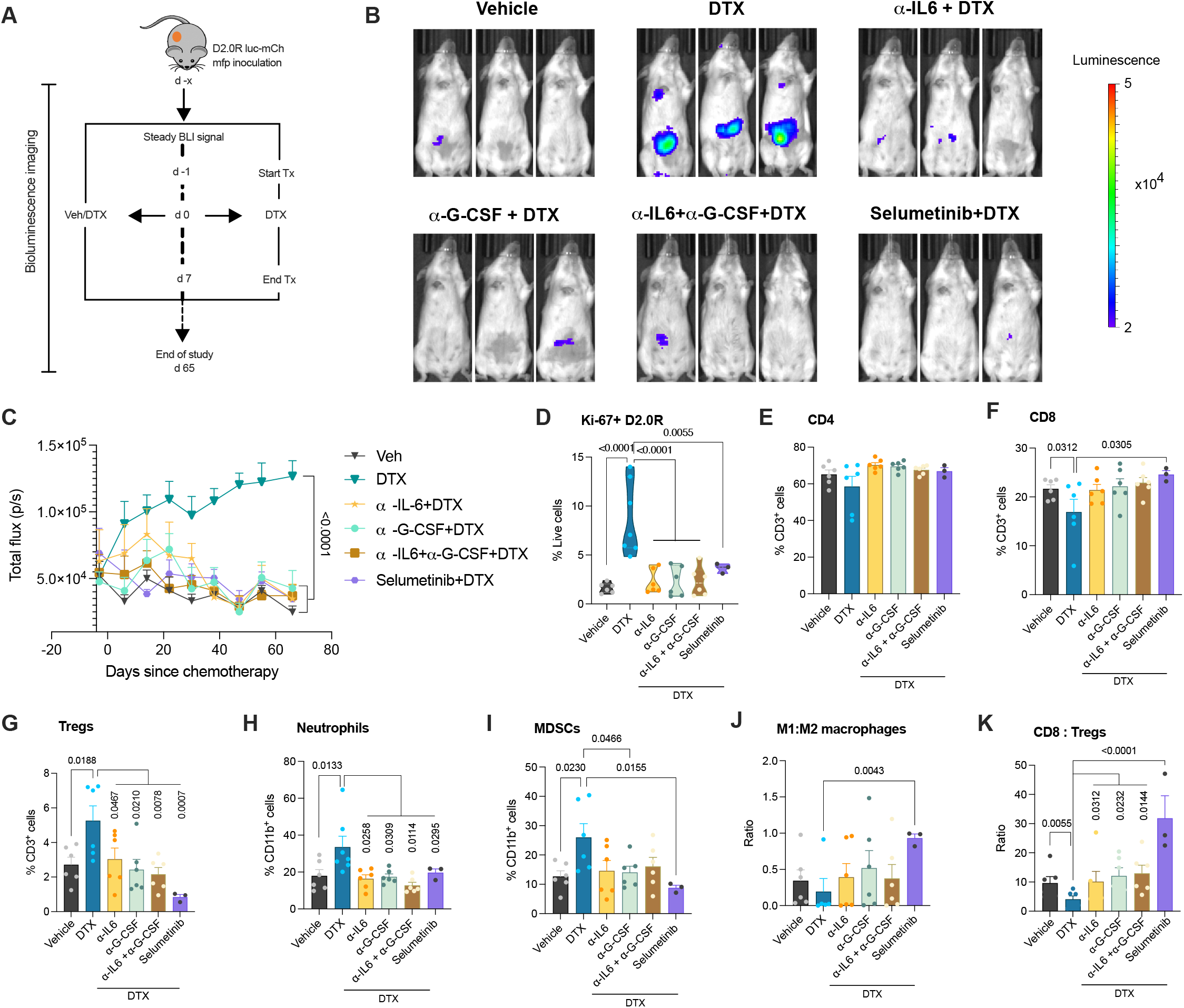
Cytokine ablation and MEK inhibition prevent chemotherapy induced breast cancer dormancy awakening in mice. (A) Schematic showing mouse model primary (mfp) dormancy by injection of D2.0R luc-mCherry, treatment groups and schedule; days since chemotherapy (d), docetaxel 8 mg/kg (DTX), neutralizing antibody or MEK inhibitor treatment (Tx) and vehicle (Veh). (B-C) Representative bioluminescence images (B) and BLI flux kinetics (C) of D2.0R luc-mCherry tumor growth in the mfp of mice in DTX (n=6), α-IL-6+DTX (n=5), α-G-CSF+DTX (n=5), α-IL-6+ α-G-CSF+DTX (n=6), selumetinib+DTX (n=6) compared with vehicle treated controls (n=6). Two-way mixed ANOVA analysis with posthoc Dunnett’s multiple comparisons test shows statistical significance between treatment groups. (D) Expression (%) of Ki67 on D2.0R luc-mCherry cells in mammary fat pad of mice from various treatment groups. One-way ANOVA measurement with posthoc Dunnett’s multiple comparisons test shows statistical significance between treatment groups. (E-I) Flow cytometric analysis showing percentage of CD4 (E), CD8 (F), Tregs (G), neutrophils (H), MDSCs (I) in the mammary fat pads/tumors of mice. Independent t-test or One-way ANOVA measurement analysis with posthoc Dunnett’s multiple comparisons test shows statistical significance between treatment groups. (J-K) Ratio of M1:M2 macrophages and CD8:Tregs in the mammary fat pads/tumors of mice. Independent t-test measurement shows statistical significance between treatment groups.

## Discussion

Despite tremendous progress made in treatments for cancer over the last two decades, cancer dormancy awakening followed by systemic recurrence continues to be a significant clinical issue. Breast cancer recurrence rates have reduced phenomenally with current treatment regimens in clinic, however there is still 6-23% chance of cancer recurrence (38), either locoregional or metastatic, within the first 5-years and about 30% rate of recurrence after 5 years from initial diagnosis and treatment (39, 40). Furthermore, the mean time from distant recurrence to death in ER negative breast cancer patients was less than three years (41). The cause of cancer recurrence varies depending on the type and size of cancer, receptor status, failure to identify local lymph node metastasis and/or undiagnosable dormant cancer cells.

Recent studies have drawn ample attention to cancer dormancy and its role in cancer recurrence (28, 42). Targeting proliferating cancer cells is the first line of thought in treating cancer, but it is important to address the silent killer within, viz dormant cancer cells. Cancer dormancy could be targeted in a number of ways: i) cancer dormancy could be prolonged eternally; ii) dormant cancer cells could be killed without being awakened from dormancy (43) or iii) killed after being awakened.

Chemotherapy is given to many cancer patients, as neoadjuvant, adjuvant or palliative therapy. Some studies have shown that chemotherapeutics induce dormancy in highly proliferating cancer cells (44), while others have shown chemotherapy induces cancer proliferation and metastasis (12, 13). Our study supports the latter view. Using a model of breast cancer dormancy, we showed for the first time that chemotherapy awakens dormant cancer cells by means of stromal injury response without affecting cancer cells directly.

We demonstrate the adverse effects of taxane-based chemotherapeutics as invoking dormant cancer outgrowth. Using a tumor stromal organoid model, we found that these deleterious effects occur in stromal cells that are sensitive to taxane killing and which release proinflammatory cytokines, such as IL-6 and G-CSF (Fig. 1-3). Clinical data from several studies show that taxane therapy increases serum IL-6 and G-CSF levels in patients (45, 46) and that higher levels of serum IL-6 is a prognostic marker for early breast cancer recurrence in patients receiving systemic therapy (47, 48). Additionally, IL-6 and G-CSF have been widely implicated in cancer invasiveness (49, 50) and recent studies have also shown a role of trans-IL-6 activating JAK/STAT signaling in dormancy awakening (51). However, the mechanistic role of IL-6 and G-CSF in inducing dormancy escape has not been investigated. By using single-cell transcriptomic and pathway analysis, we found that IL-6 and G-CSF engagement in non-canonical signaling via the ERK/MAPK pathway, involved in fibrosis and wound healing response (52, 53), to be crucial in cancer cells outgrowth (Fig. 3), instead of the canonical JAK/STAT signaling pathway involved in inflammatory response as shown by previous studies (17, 51). The gene signatures identified in the chemotherapy group agreed with previous studies on dormant-emergent cancer and chemoresistance (18, 42). Furthermore, using a syngeneic mouse model of breast cancer dormancy, both orthotopic and metastatic, we confirmed the deleterious effects of taxane-based chemotherapy in dormancy awakening *in vivo*, and systemic release of proinflammatory cytokines IL-6 and G-CSF. Using single-cell transcriptomics analysis we confirmed that the source of IL-6 and G-CSF was indeed fibroblasts/CAFs and fibrocytes, in agreement with our *in vitro* data and previous findings showing stromal IL-6 plays a role in invoking cancer cell proliferation (11). Based on CellChat analysis we found that upon docetaxel treatment, fibroblasts/CAFs have the strongest and highest number of interactions with other cell types in the tumor. We also found that in the docetaxel-treated mice tumor IL-6 signaling pathway was strongly signaling between fibroblasts/CAFs and endothelial cells with cancer cells. Interestingly, we also found that *in vivo* G-CSF/CSF3 signaling was primarily between fibrocytes and neutrophils or M2 macrophages suggesting robust activation of these suppressive immune cells in the cancer microenvironment, data which was corroborated by increased expression of suppressive cytokine and chemokine genes (Fig. 6) strongly suggesting a role of M2 TAMs in chemotherapy induced pro-tumor signaling (54).

Upon investigating the mammary tumor tissue from docetaxel treated mice, we found an increase in pro-tumor immune infiltrates including M2 macrophages, MDSCs, Tregs, γδT cells and an enriched gene signature comprising *Vim, Spp1, S100a6, Ccl2* and *Lgals1* genes. Previous studies have shown that *Vim, Spp1* and *Ccl2* are IL-6 responsive genes (55-57). Based on Cellchat analysis we found that Spp1 signaling was initiated by cancer cells in networking with other cells in the tumor microenvironment (TME). Importantly, *in vivo* blockade of IL-6 or G-CSF signaling or MEK signaling prevents DTX induced dormancy escape. Taken together, our findings point to a single signaling mechanism (MEK signaling) by which DTX chemotherapy causes breast cancer dormancy outgrowth (*in vitro* and *in vivo*).

In the *in vivo* model of breast cancer, we noticed that dormant cancer cells were awakened with just a single dose of chemotherapy. Following up on this, we were unable to target the awakened/proliferating cells with subsequent cycles of chemotherapy (Fig. 4 and Fig. S3). Later, we confirmed by transcriptomic analysis that chemotherapy mediated cancer dormancy escape induced several survival cues in cancer cells including chemoresistance, EMT, invasiveness and inflammatory phenotype, evident from the expression of *Cav1, Col3a1, Col5a1, Spp1, Fth1, Hmox1, Il6, Cxcl10, Cxcl1, Ccl2*, etc. Interestingly, our findings showed that chemotherapy not only awakened dormant cancer cells, but also caused clonal propagation of cancer cells, wherein one cluster gained chemoresistance phenotype expressing genes such as *Cav1, Col3a1* and *Col5a (58, 59)*, while the other gained an inflammatory phenotype expressing genes (60, 61) such as *Cxcl1, Cxcl10, Ccl2* (Fig. S5G).

There are some caveats to our study. First, the organoid system contains only cancer cells and stromal cells (i.e., fibroblasts and endothelial cells). However, the *in vivo* mammary tumor microenvironment revealed several immune cells being recruited upon stromal injury to play protumor roles in facilitating tumor growth and progression (Fig. 5-6). Secondly, even though our study revealed a role of taxane chemotherapy in stromal injury mediated dormancy awakening *in vitro* and *in vivo*, nonetheless our model does not address the complexity and heterogeneity of the different populations of cancer cells and the microenvironments within a complex tumor.

In conclusion, our study has identified key secretory factors (IL-6 and G-CSF) as mediators of taxane induced dormancy awakening in a breast cancer model (Fig. 8). Additional preclinical studies in other tumor types and with other chemotherapies are warranted as are studies in the clinic to monitor changes in the levels of cytokines, chemokines, and growth factors in patients before and after chemotherapy. Our studies suggest that administration of IL-6 or G-CSF signaling inhibitors in the peri-chemotherapy period (at least in patients who might be expected to have a surge of IL-6 or G-CSF levels due to chemotherapy), might decrease tumor recurrence and improve survival outcomes.

**Figure 8.**
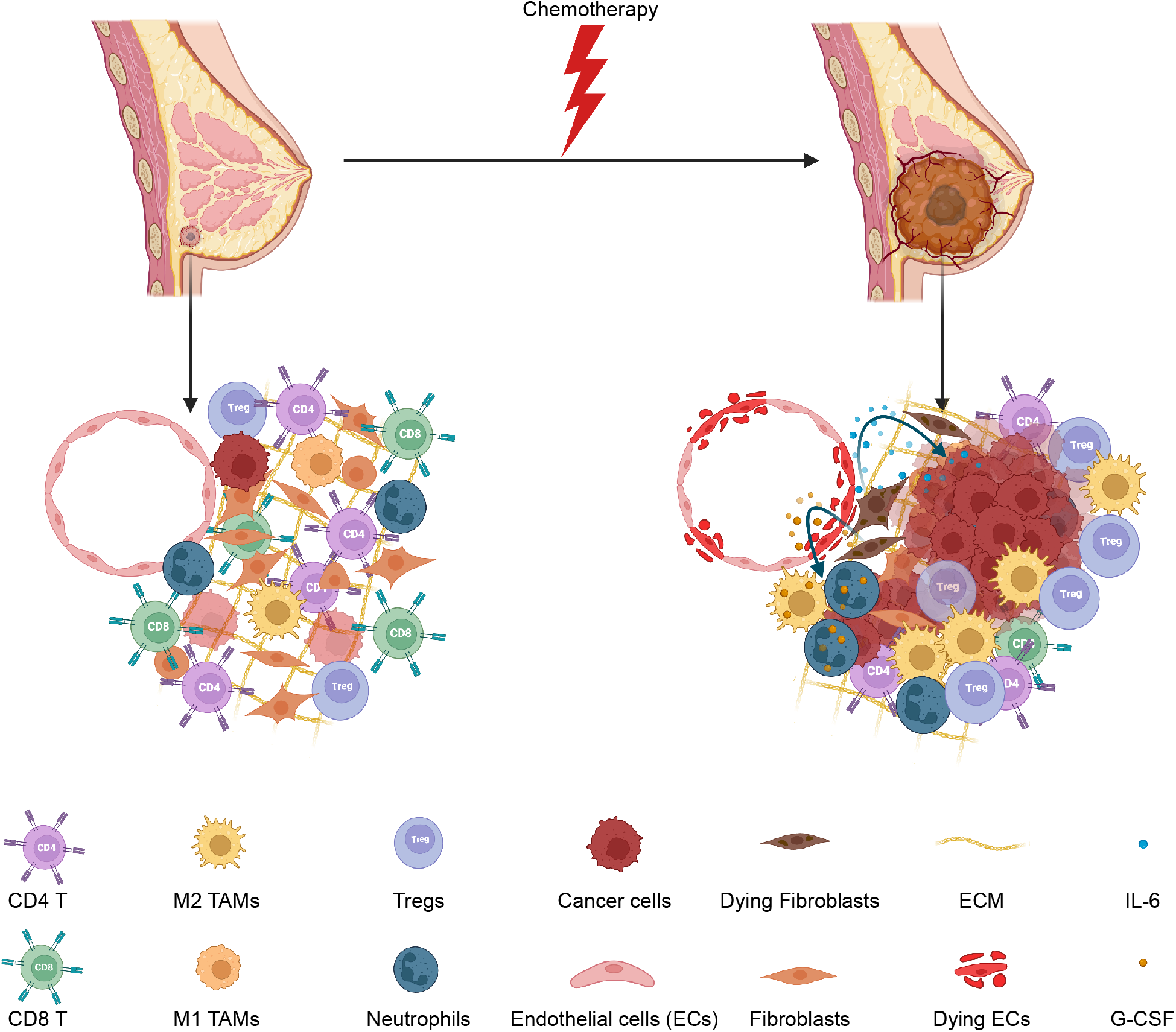
Summary schematic of chemotherapy induced stromal injury and dormancy outgrowth. This schematic shows a dormant tumor in the breast (on the left) with very few dormant cancer cells, healthy stromal cells, and an anti-tumor immune microenvironment. Chemotherapy injures stromal cells (shown by dying stromal cells) releasing IL-6 and G-CSF, which in turn awaken dormant cancer cells, recruit pro-tumor neutrophils and M2 macrophages. Furthermore, more protumor immune infiltrates such as Tregs and fewer anti-tumor CD8 T cells were seen after dormancy outgrowth, confirming an overall pro-tumor microenvironment facilitating tumor growth.

## Acknowledgments

We thank Dr. Sudarshan Malla and Rebecca Pankove for their input in the preliminary stages of this study. We thank Dr. Dmitry Shayakhmetov for allowing us to use Incucyte and ChemiDoc XRS+.

Experiments in Dr. Manoj Bhasin and Dr. Swati Sharma Bhasin’s laboratory were supported by Emory lab startup funds to Dr. Bhasin.

## Author Contributions

R.G. conceived, designed the study, carried out experiments and data analysis, prepared figures and wrote the manuscript. V.P.S. conceived and supervised research, contributed to manuscript preparation and critically edited the manuscript. S.S.B. and M.B. contributed to manuscript preparation, supervised transcriptomics studies, and critically edited the manuscript. B.E.T. performed single cell RNA library preparation and sequencing and U.K. performed bioinformatics analysis of transcriptomics data and contributed to manuscript and figure preparation. N.R.C contributed to data analysis using IPA and manuscript preparation.

