## Supplemental Figure S1 for "Taxane chemotherapy leads to breast cancer dormancy escape by stromal injury mediated IL-6/MAP2K signaling"

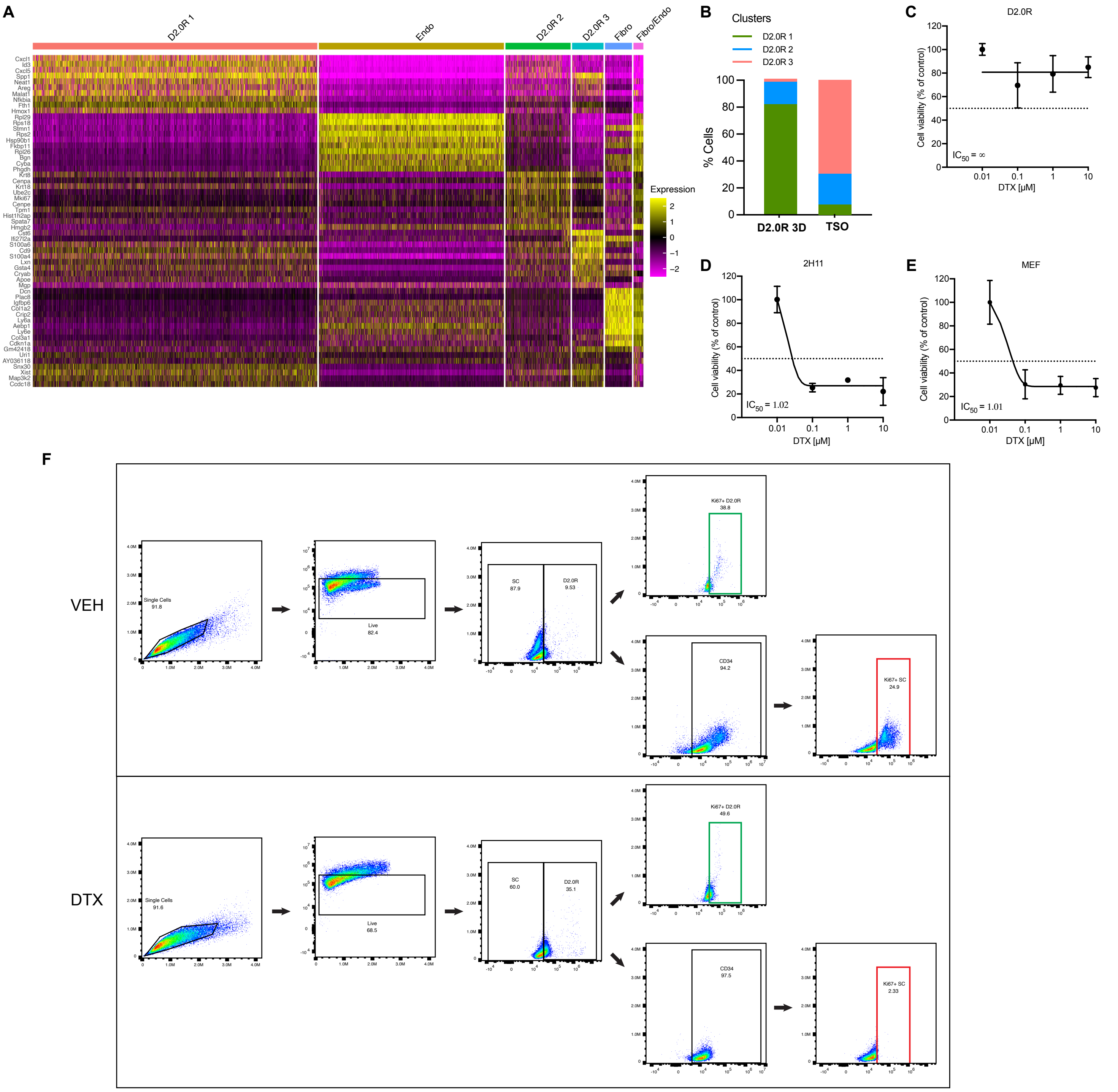


**Fig S1. Cell viability assay and flow gating in vehicle and docetaxel treatment.** (A) Heatmap showing the top 10 DEGs in each cluster in a merged dataset. (B) Percentage of the different clusters of cancer cells from D2.0R 3D and TSO, per total cancer cells in the respective datasets. (C-E) Dose response curves of cancer cells (D2.0R) (n=5), endothelial (2H11) (n=4) and fibroblasts (MEF) (n=3) to varying concentrations of DTX (0-10 µM). (F) Representative images of flow gating strategy for singlets, live cells and Ki67+ D2.0Rs (mCherry) (green box) or Ki-67+ stromal cells (2H11:MEF) (red box) in VEH and DTX treated tumor stromal organoids.
