## Supplemental Table 1 for "Taxane chemotherapy leads to breast cancer dormancy escape by stromal injury mediated IL-6/MAP2K signaling"

**Table S1. Dormancy associated genes upregulated in cancer cells in TSO relative to D2.0R 3D**

| Genes | p_val_adj | avg_log2FC |
| --- | --- | --- |
| *Cst6* | 2.01E-104 | 2.15954264 |
| *Mgp* | 1.02E-13 | 1.54297919 |
| *Mme* | 4.00E-14 | 0.54430503 |
| *Thbs1* | 3.71E-11 | 0.53548017 |
| *Gas6* | 1.44E-05 | 0.48884647 |
| *Timp2* | 3.49E-20 | 0.47096683 |
| *Ndrg1* | 4.05E-06 | 0.34776753 |
