## Supplemental Figure S2 for "Taxane chemotherapy leads to breast cancer dormancy escape by stromal injury mediated IL-6/MAP2K signaling"

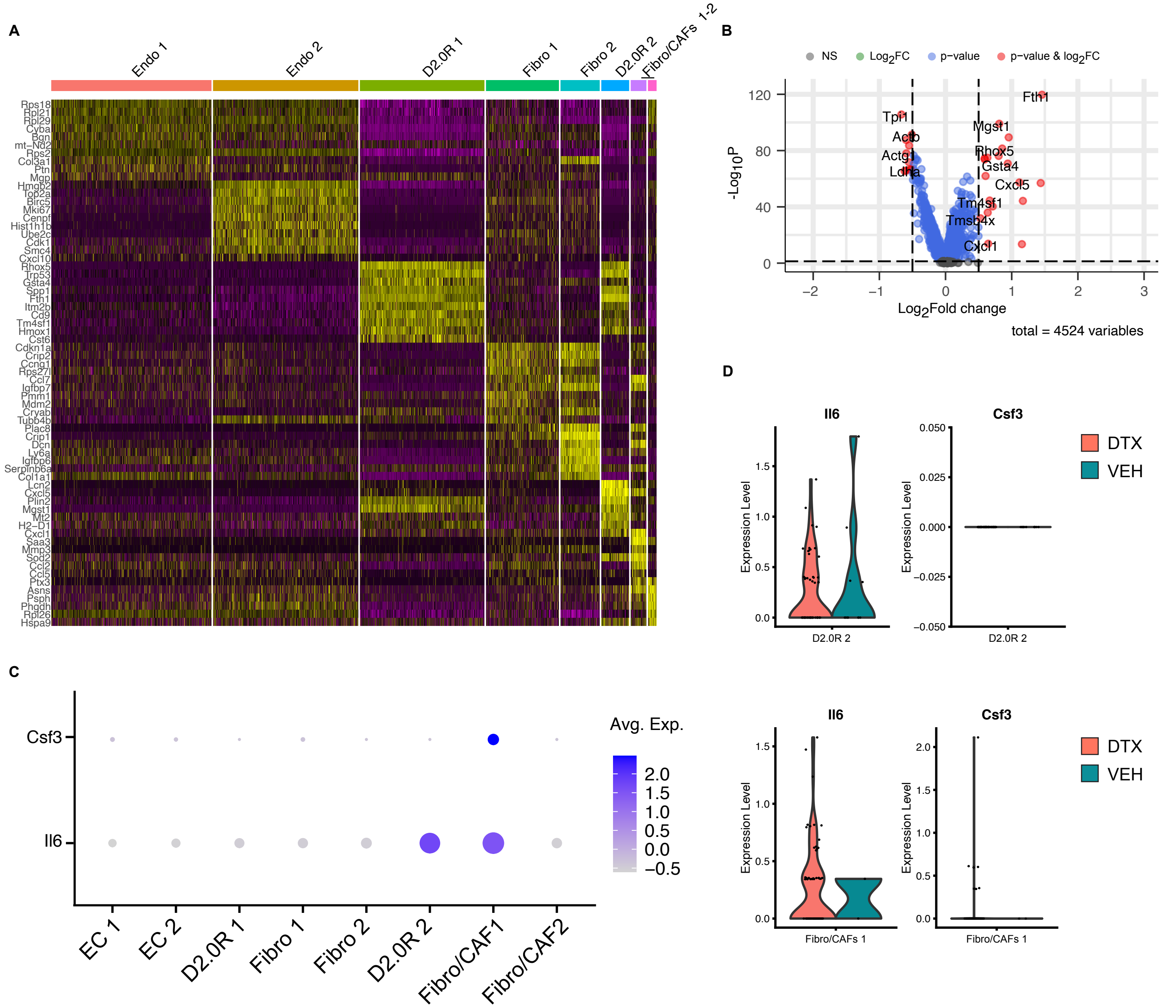


**Fig S2. Transcriptomics data DEGs in the clusters and upon docetaxel treatment.** (A) Heatmap showing the top 10 DEGs in each cluster in a merged dataset. (B) Volcano plot showing significantly differentially expressed protein-coding genes in single cell suspension of tumor stromal organoids based on scRNA-seq data from docetaxel treated compared with vehicle treated controls. Transcripts with FC > 0.5 and adjusted P value < 0.05 are highlighted in red. (C) Dot plots of Il6 and Csf3 genes from merged samples. Dot size indicates the proportion of cells in each cluster expressing a gene and color shading indicates the relative level of gene expression. (D) Violin plots showing Il6 and Csf3 gene expression levels in D2.0R 2 cancer cell cluster (top) and Fibro/CAF1stromal cell cluster (bottom).
