## Supplemental Figure S3 for "Taxane chemotherapy leads to breast cancer dormancy escape by stromal injury mediated IL-6/MAP2K signaling"

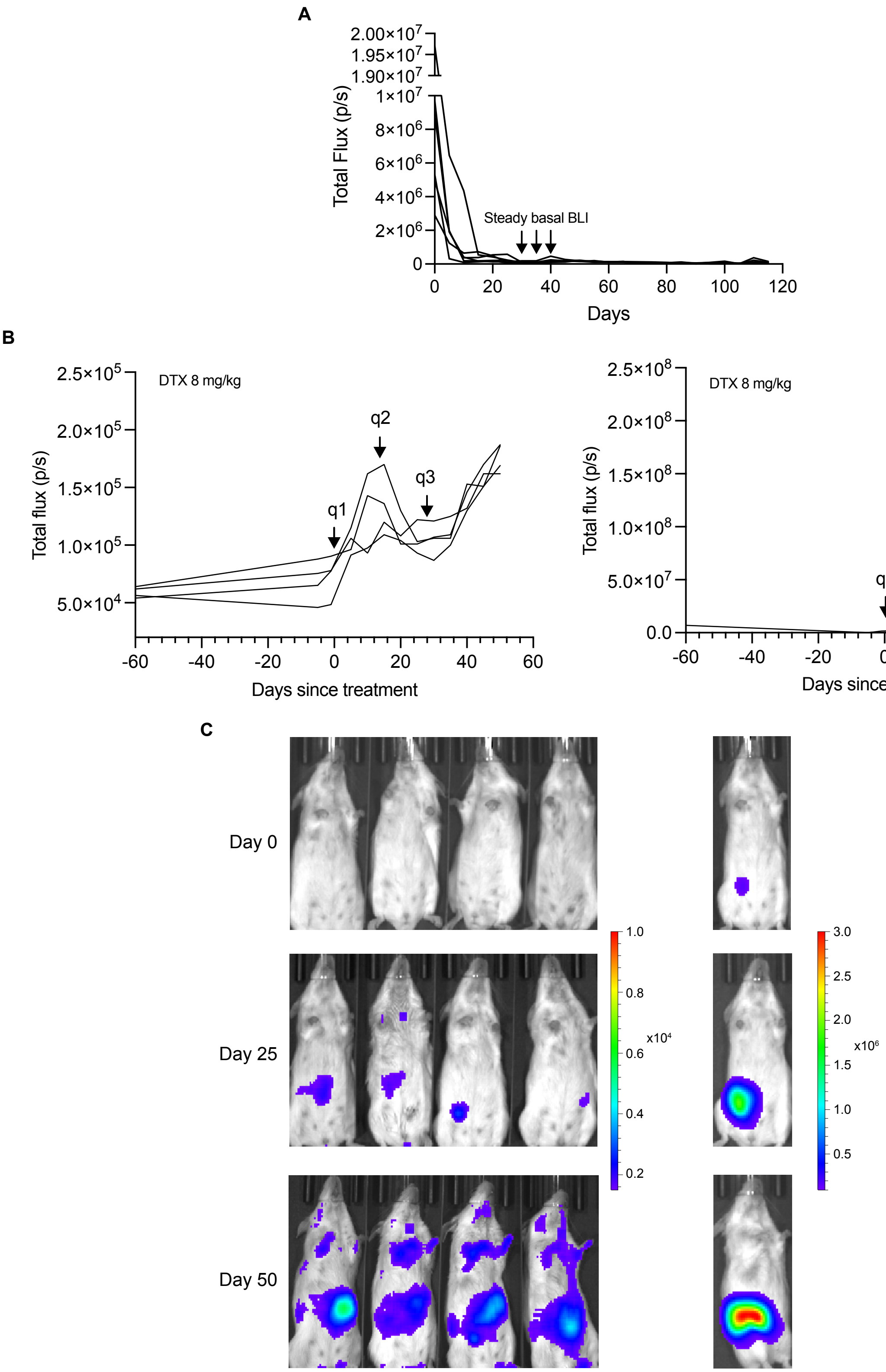


**Fig S3. Awakened cancer cells did not succumb to repeated cycles of chemotherapy.** (A) Representative BLI flux kinetics of D2.0R luc-mCherry tumor growth in the mfp of untreated mice followed for ~4 months (n=6). (B-C) Representative BLI flux kinetics (B) and bioluminescence images (C) of D2.0R luc-mCherry tumor growth in the mfp of mice treated with 3 cycles of docetaxel (n=5).
