## Supplemental Figure S4 for "Taxane chemotherapy leads to breast cancer dormancy escape by stromal injury mediated IL-6/MAP2K signaling"

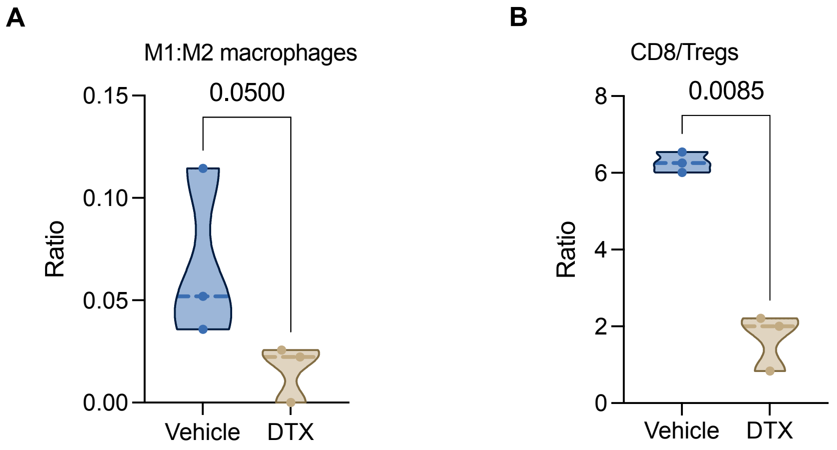


**Fig S4. Tumor immunosuppressive signature upon docetaxel treatment.** (A-B) Ratio of M1:M2 macrophages (A) and CD8:Tregs (B) in the mammary fat pads/tumors of mice treated with vehicle or docetaxel (n=3, each). Independent t-test measurement shows statistical significance between treatment groups.
