## Supplemental Figure S5 for "Taxane chemotherapy leads to breast cancer dormancy escape by stromal injury mediated IL-6/MAP2K signaling"

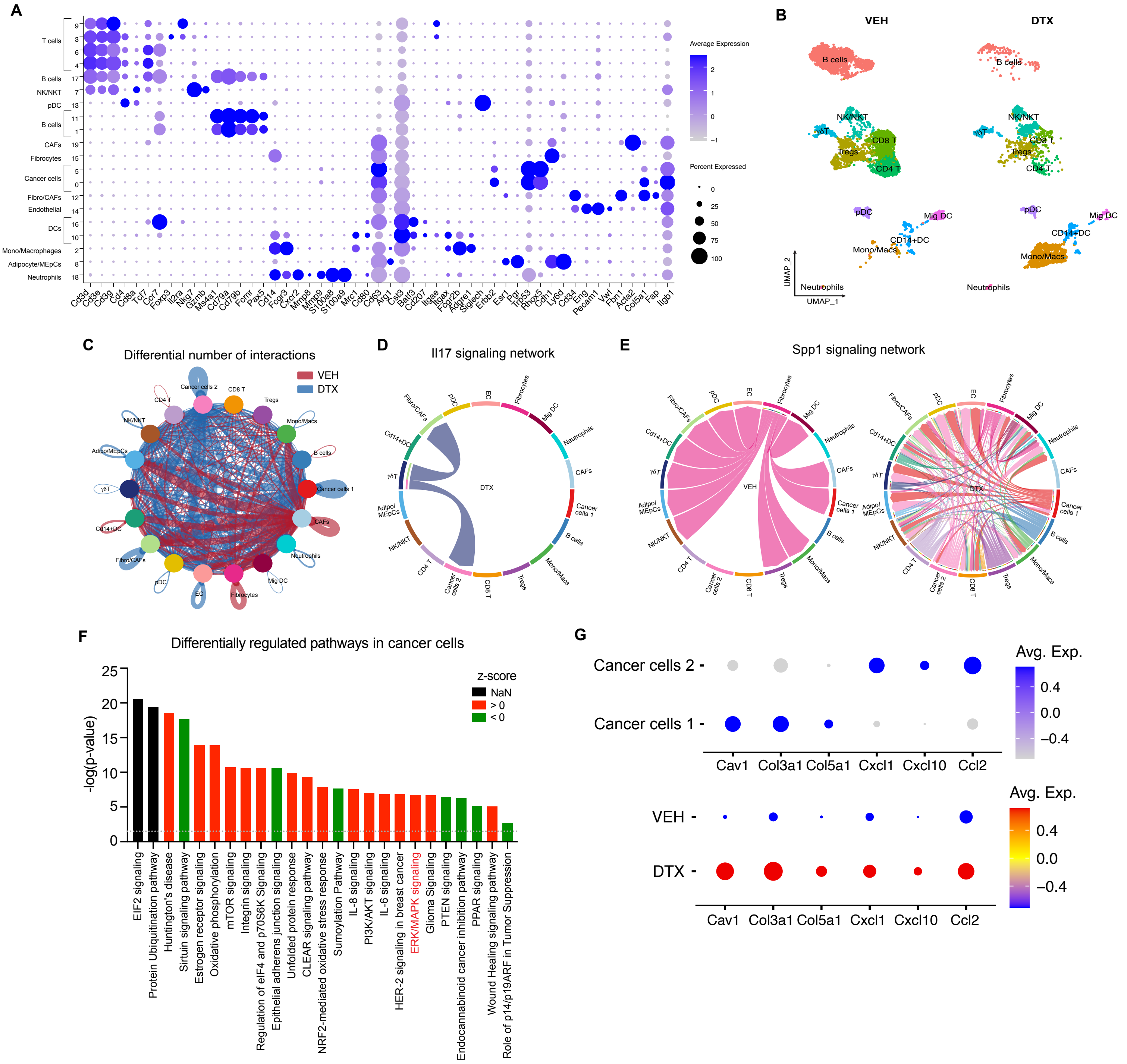


**Fig S5. Transcriptomics analysis- cluster markers, cell-cell communication, and cancer cell phenotypes.** (A) Cell type identifying cluster markers with clusters grouped by major phenotype classifying genes. (B) UMAP presentation of major immune cell types and associated clusters in vehicle (VEH) and docetaxel (DTX) treated murine mammary fat pad/tumor (n=2, each treatment group). (C) Connectivity map: circles represent source and target clusters, respectively; Gradience in the color shows lowest (light) to highest (dark) number of network connections; red gradient- Vehicle and blue gradient- DTX. (D) Chord connections between cell types shows *Il17* signaling network in DTX dataset. (E) Chord connections between cell types shows *Spp1* signaling network in VEH and DTX datasets, respectively. (F) Differentially regulated ingenuity pathways enriched or downregulated upon docetaxel treatment compared with vehicle controls in cancer cells in the mfp/tumor based on differentially expressed RNAs. -log (p-value) for each pathway are indicated on the x-axis; Color red or green indicates positive or negative z-score, respectively. (G) Dot plots of selected markers in cancer cells 1 and cancer cells 2 from merged dataset (top) and grouped by treatment, VEH and DTX (bottom). Dot size indicates the proportion of cells in each cluster expressing a gene and color shading indicates the relative level of gene expression.
